# Microtubule dynamics are defined by conformations and stability of clustered protofilaments

**DOI:** 10.1101/2024.11.04.621893

**Authors:** Maksim Kalutskii, Helmut Grubmüller, Vladimir A Volkov, Maxim Igaev

## Abstract

Microtubules are dynamic cytoskeletal polymers that add and lose tubulin dimers at their ends. Microtubule growth, shortening and transitions between them are linked to GTP hydrolysis. Recent evidence suggests that flexible tubulin protofilaments at microtubule ends adopt a variety of shapes, complicating structural analysis using conventional techniques. Therefore, the link between GTP hydrolysis, protofilament structure and microtubule polymerization state is poorly understood. Here, we investigate the conformational dynamics of microtubule ends using coarse-grained modeling supported by atomistic simulations and cryo-electron tomography. We show that individual bent protofilaments organize in clusters, transient precursors to a straight microtubule lattice, with GTP-bound ends showing elevated and more persistent cluster formation. Differences in the mechanical properties of GTP- and GDP-protofilaments result in differences in intra-cluster tension, determining both clustering propensity and protofilament length. We propose that conformational selection at microtubule ends favors long-lived clusters of short GTP-protofilaments that are more prone to form a straight microtubule lattice and accommodate new tubulin dimers. Conversely, microtubule ends trapped in states with unevenly long and stiff GDP-protofilaments are more prone to shortening. We conclude that protofilament clustering is the key phenomenon that links the hydrolysis state of single tubulins to the polymerization state of the entire microtubule.

## Introduction

Microtubules are conserved cytoskeletal polymers which play crucial roles in processes ranging from cell division to neuronal homeostasis.^1–3^ Moreover, microtubule assembly and disassembly can produce mechanical forces in the piconewton range.^4–8^ The remarkable property of microtubules to stochastically transition between phases of growth and shortening^9^ has been linked to the binding and hydrolysis of GTP coupled to the addition of tubulin to microtubule ends.^10,11^ The idea that GTP is required for polymerization, but GTP hydrolysis upon polymerization renders microtubules intrinsically unstable is largely undisputed and is reflected in the concept of a stabilizing “GTP cap”.^9,12^ The GTP cap at the microtubule end is proposed to span hundreds of nanometers of microtubule length^13–15^ and to maintain polymer stability, while its loss can trigger depolymerization.

While the GTP cap is a simple, yet fundamental concept for understanding tubulin polymerization, it does not explain the coupling between GTP hydrolysis and microtubule (dis)assembly, limiting predictions of microtubule behavior in various physiological and pathological contexts. The first models describing the mechanochemical cycle of tubulin^9,12^ were proposed before the crystal structure of tubulin was resolved.^16,17^ Early electron microscopy studies revealed growing microtubule ends as mostly blunt or carrying sheet-like extensions, and shortening ones as flared, with protofilaments curling outwards.^18–21^ Despite the fact that these studies relied on 2D projections and did not resolve 3D structures of microtubule ends explicitly, they elegantly explained why GDP-bound tubulin would not polymerize: Only GTP-bound tubulin can adopt the necessary straight(er) conformation to bond with neighboring tubulins. However, subsequent structural characterization found no significant differences between the conformations of GTP- and GDP-bound tubulin in solution,^22–24^ in crystals^25–28^ and in silico,^29–32^ challenging this hypothesis. Despite this controversy, the idea of nucleotide-dependent tubulin curvature has been incorporated into several minimal models of microtubule assembly.^33–39^

Recent cryo-electron tomography (cryoET) work by McIntosh, Gudimchuk et al.^40,41^ and others^42^ resolved 3D structures of both growing and shortening microtubule ends explicitly, and showed curled protofilaments in both polymerization states, in vitro as well as in vivo, implying profound changes to the tubulin polymerization paradigm. Moreover, cryoET reconstructions revealed that protofilaments were flexible in the radial plane of bending, whereas stepping out of that plane was less than 10% in terms of protofilament length, suggesting a much higher tangential rigidity and no interactions between adjacent protofilaments.^40,41^ Consequently, it was proposed that microtubule growth is achieved by thermal fluctuations of similarly curved and independent protofilaments, with only the bonds between GTP-tubulins being stable enough to maintain a straight lattice. This idea was further explored by the same authors using Brownian Dynamics modeling.^40,41,43^

Most recently, we and others have performed large-scale atomistic simulations of complete GTP- and GDP-microtubule end models over the course of microseconds.^44–46^ In particular, our study^45^ confirmed that all microtubule ends tend to be flared regardless of the nucleotide state; however, other key observations were incompatible with the original findings.^40–42^ Specifically, protofilaments were flexible both within and outside the radial plane, and clusters of laterally connected protofilaments were directly observed as the system was minimizing the mechanical frustration during the relaxation. The nucleotide state affected this delicate balance by modulating both the tangential flexibility of individual protofilaments and the energetics of lateral interactions. We hence predicted that protofilament clusters might be important structural intermediates that lower the activation barrier for the formation of a straight microtubule lattice. We further hypothesized that kinetic control over cluster formation might be a key determinant of the self-assembly mechanism and dynamic instability. However, the computational cost of current atomistic simulations did not allow us to observe reversible association and dissociation of protofilaments into clusters and, therefore, to explore how this would guide the time evolution of a microtubule end at much longer timescales. As a result, there is still no consensus about the true conformational ensemble of the microtubule end and the thermodynamic and kinetic determinants of its capacity to elongate.

But why is it difficult to reach a consensus? Our understanding of the microtubule end dynamics is currently limited by two main challenges. On the experimental side, the transient nature of microtubule ends hinders real-time, high-resolution measurements. Whereas single-particle cryo-electron microscopy offers near-atomic static snapshots of microtubule segments away from the end,^15,47–51^ variable structures of protofilaments at microtubule ends cannot be resolved due to the heterogeneity of protofilament shapes. In turn, cryoET can resolve the structure of microtubule ends without averaging,^40–42,52^ but with a much lower signal-to-noise ratio, rendering the structural analysis at the level of tubulin-tubulin interactions challenging. Furthermore, fluorescence microscopy can track microtubule ends in real time but lacks spatial resolution to provide structural information.^53–56^ On the theoretical side, multiscale computational approaches to predict the impact of the bound nucleotide on the dynamics of microtubule ends are not available, while accurate atomistic simulations studying large-scale processes such as fluctuations of the microtubule end are too computationally expensive to cover the relevant timescales. In addition, existing minimalistic models often tend to oversimplify microtubule structure and dynamics, thus limiting quantitative studies. For example, even the most advanced models^41,43^ miss the complex bending-torsional dynamics of protofilaments as well as important correlations caused by intermolecular interactions.^44,45,57^ To obtain a quantitative understanding of structure-dynamics relationships in microtubule assembly, new integrative strategies are therefore required.

In this work, we examine the structure and dynamics of microtubule ends in both nucleotide states at millisecond timescales using coarse-grained (CG) modeling informed and parametrized by atomistic simulations. To this end, an ab initio approach is used to construct an elastic CG model of microtubule end dynamics accounting for both the bending-torsional elasticity of individual protofilaments and the correlations caused by neighbor interactions. We compare results of these simulations to experimental structures of microtubule ends determined using a combination of cryoET and deep-learning-based image denoising, allowing us to increase the precision of segmentation used to determine 3D coordinates of individual tubulin monomers within a flared microtubule end. CG simulations and cryoET reveal that growing microtubule ends feature longer-lived clusters involving a larger number of protofilaments, as compared with the shortening ends. Our modeling also predicts that excess tensile stress in the clusters leads to irreversible protofilament ruptures and tubulin dissociation. Moreover, the rate of these protofilament ruptures is elevated in GTP-state, explaining why growing microtubule ends have shorter and more uniform protofilaments, in agreement with our cryoET measurements. In contrast, GDP-microtubule ends shorten because they get trapped in states with long uneven protofilaments or pairs thereof, thus increasing the free energy barrier to a straight microtubule lattice.

## Results and Discussion

### Protofilament clusters are present at microtubule ends in both polymerization states

Our previous all-atom molecular dynamics (MD) study^45^ showed that the non-equilibrium relaxation of the microtubule end structure occurs at microsecond time scales and is driven by a “tug-of-war” between the bending-torsional elasticity of protofilaments and lateral interactions between them (Fig. 1A). However, neither was it possible to converge these computationally expensive simulations to a steady state in which the processes of protofilament clustering and separation would be in equilibrium, nor were experimental structural data supporting this hypothesis available. Here, we ask whether protofilament clusters generally exist at the ends of growing and shrinking microtubules and, if so, whether a model can be derived to accurately describe and understand this phenomenon from fundamental principles.

**Figure 1.**
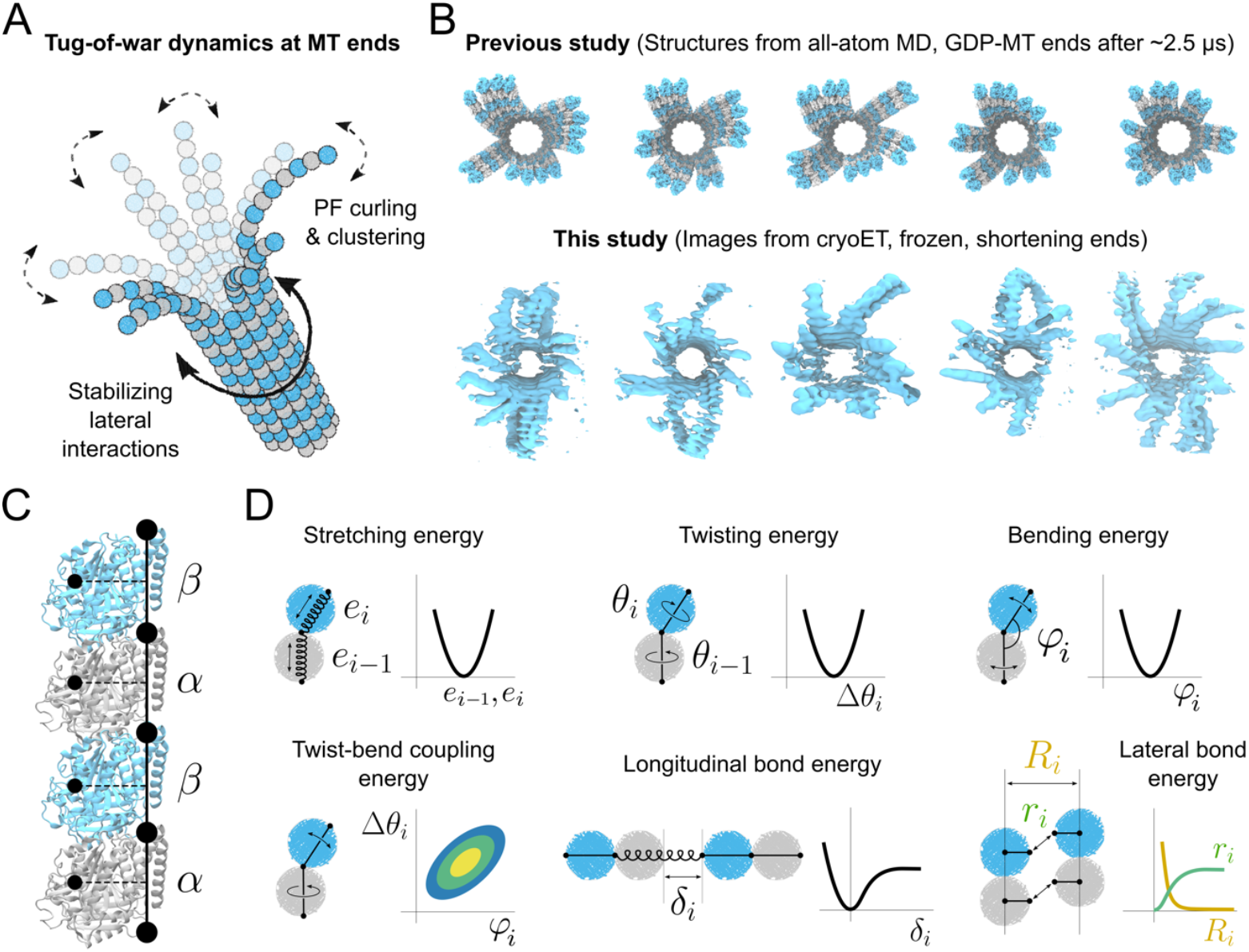
**(A)** Cartoon illustration of the “tug-of-war” principle: the curved shape of protofilaments at the microtubule end is geometrically incompatible with the straight lattice, resulting in intermediate clusters of partially straightened protofilaments. **(B)** Comparison of the simulated GDP-microtubule ends from our previous study^45^ (top) with exemplary 3D rendered volumes of shortening microtubule plus-ends obtained in this study (bottom). **(C)** CG representation of a protofilament (black circles with solid and dashed lines) mapped onto its atomistic structure. *α*- and *β*-tubulin are shown as gray and cyan ribbons, respectively. **(D)** Minimal elastic coarse-grained model of a microtubule end. Each protofilament is modeled as a set of nodes connected by stretchable and twistable springs. Coupling between bending and twisting of neighboring strings is introduced to better reproduce the atomistic dynamics. All deformations are described by harmonic potentials except those between individual tubulins.

To resolve the structures of microtubule ends in growing and shortening states, we performed cryoET on samples containing dynamic microtubules polymerized from purified bovine brain tubulin (see Materials and Methods). Cryo-CARE denoising allowed us to reduce high-frequency noise sufficiently enough to resolve individual flaring protofilaments at microtubule plus-ends in 3D.^58^ Since the microtubules were polymerized from GMPCPP seeds, the majority of them had 14 protofilaments, which enabled an unambiguous comparison of the reconstructed tomograms with our all-atom MD simulations of 14-protofilament microtubule ends. Figure 1B shows the end-on view of protofilament flares of the GDP-microtubule plus-ends simulated for ∼2.5 μs from our previous study and some exemplary 3D tomograms of the shortening microtubule plus-ends obtained in this study (see Fig. S1A for the comparison of growing microtubule ends).

Even without further analysis of the tomograms, which will be presented in detail below, protofilament clusters were clearly detectable as the resolution was sufficient to observe individual tubulin molecules. Contrary to previous reports,^40,41^ the protofilaments in our samples deviated from their radial planes to form clusters with their neighbors – an observation which we attribute to the improved denoising procedure. Our previous MD simulations (Fig. 1B, top row) already established that a soft tangential mode of protofilament motion was responsible for the out-of-plane deviations (“tangential swing”; see Movie S3 in here^45^). Furthermore, approximately 84% of the reconstructed microtubule tomograms showed a global left-handed twist pattern, i.e. the protofilaments twist-bent counterclockwise in the direction of microtubule growth. This pattern – also clearly observed in the simulated microtubule ends – likely resulted from the torsional component in the protofilaments’ main bending mode (see Movie S3 in here^45^), which caused asymmetric exploration of the conformational space at the microtubule end. Together, these observations lead us to conclude that protofilament clusters are not an artifact of cryoET reconstruction or simulation, but rather structural intermediates characteristic of both microtubule polymerization states.

### A coarse-grained model allows access to sub-millisecond dynamics of the microtubule end

We constructed an elastic CG model to quantify the dynamics and energetics of protofilament clustering at the microtubule end. To account for the radial and tangential elasticity of protofilaments, each tubulin dimer was represented by three CG beads connected by stretchable and rotatable springs (Fig. 1C; see Materials and Methods, Fig. S1B and Table S1 for a detailed description of the model geometry). Thus, the black dots in Fig. 1C represent sites at which model parameters are relevant, while the minimal simulated entity in our model is a tubulin dimer. This type of bead assignment reflects the fact that protofilaments bend and twist at “hinge” regions located at the intra- or inter-dimer interfaces between α- and β-subunits.^45,57,59^ Therefore, unlike in many previous models, a CG bead in our model does not coincide with a tubulin monomer but instead is shared by two neighboring monomers except when it is a terminal one. Furthermore, all elementary deformations in the model were designed to be harmonic, and the corresponding mechanical parameters for every triplet of CG beads were set to depend only on the nucleotide state and on whether it described an intra- or inter-dimer interface. To describe protofilament-protofilament interactions and to allow for tubulin dissociation, we also introduced breakable lateral and longitudinal bonds. Figure 1D schematically summarizes the elementary strains, and the graphs beside them show the potential function under consideration. The model parameters were derived from our previous^45^ and newly produced all-atom MD simulations (see Materials and Methods, Fig. S1C–E and Table S2 for a detailed description of the parametrization procedure).

### Nucleotide state modulates “tug-of-war” dynamics by controlling protofilament cluster size, number and stability

The model introduced above allowed us to predict the stochastic time evolution of the microtubule end at sub-millisecond timescales currently inaccessible to both high-resolution microscopy and all-atom MD simulations. It further enabled direct comparison to the structures obtained via cryoET (which are presented in detail below). In our previous MD study,^45^ we predicted that the size and stability of intermediate protofilament clusters should determine the probability of growth or shortening. Consequently, larger and more stable clusters should increase the growth probability without significantly changing the overall flared shape of the microtubule end. To test this prediction, we carried out Brownian Dynamics simulations of our model to obtain conformational ensembles of microtubule ends with 6 dimers per protofilament as a function of the nucleotide state and the lateral interaction energy. For each pair of these parameters, approximately 30 × 200 μs of CG trajectories were generated (see Materials and Methods for details on simulations and analysis and Movies S1 and S2 for exemplary trajectories of the GTP- and GDP-microtubule end dynamics).

We first quantified how many clusters were formed and what fraction of protofilaments participated in clustering for both nucleotide states. As expected, in the absence of lateral interactions (*U*_*lat*_ = 0 kJ/mol, where *U*_*lat*_ is the attractive part of the lateral interaction potential in Fig. 1D), the protofilaments did not engage in lateral interactions. With increasing *U*_*lat*_, an increasing number of protofilament clusters emerged (Fig. 2A) that also gradually grew in size (Fig. 2B). For very strong lateral interactions *U*_*lat*_ > 50 kJ/mol, the clustering was limited by the number of available protofilaments in the microtubule (14 in our simulations), reflected in a slight drop in the average number of clusters.

**Figure 2.**
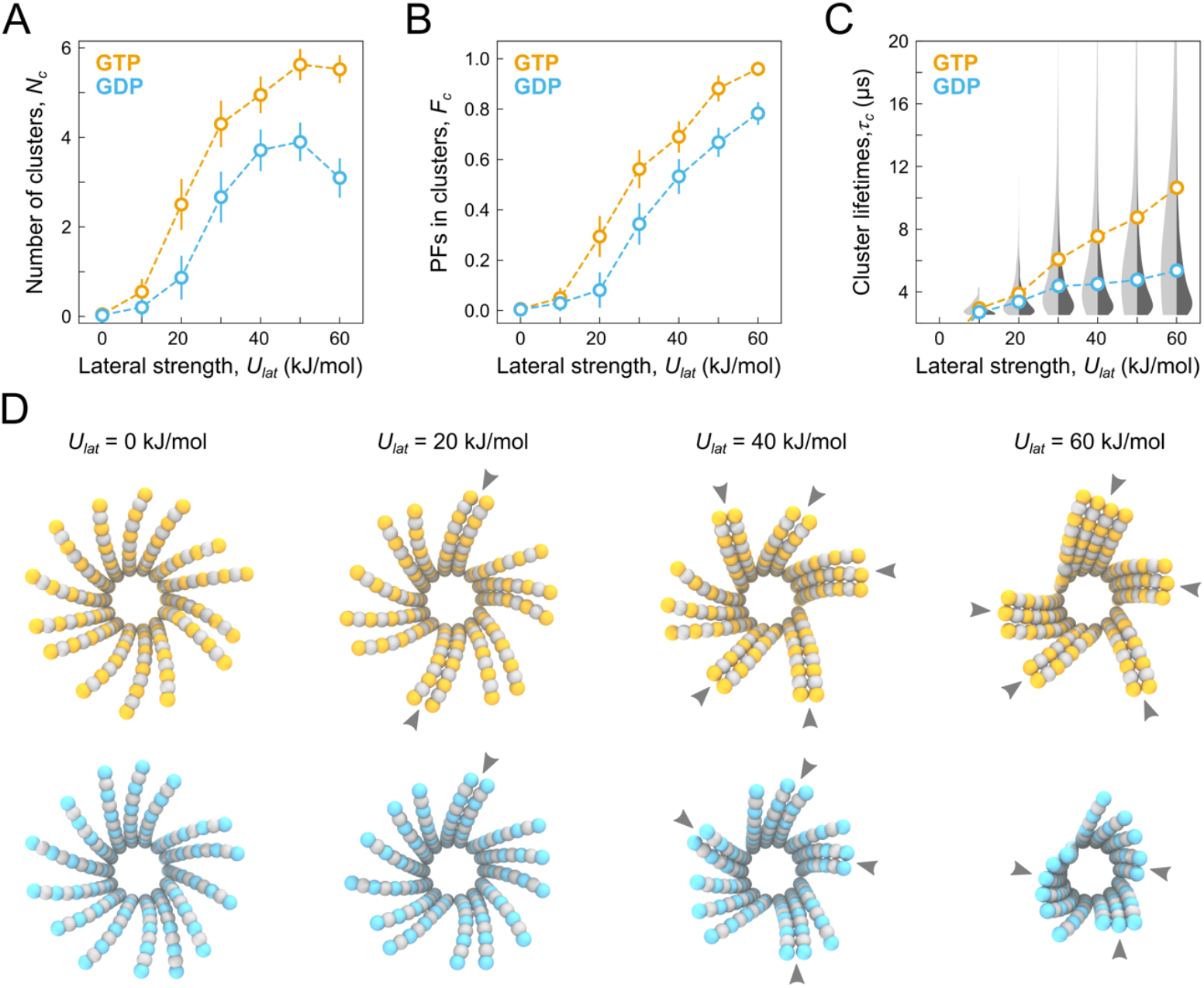
**(A)** The average number of clusters, *N*_*c*_, **(B)** the average fraction of protofilaments in clusters, *F*_*c*_, and **(C)** the average cluster lifetime, *τ*_*c*_, plotted as a function of nucleotide state (orange for GTP and cyan for GDP) and lateral interaction energy, *U*_*lat*_. For *Nc* and *F*_*c*_, the error bars indicate standard deviations calculated over all time frames (*n*=8000) and all simulation replicas (*m*=30). For *τ*_*c*_, no error bars are provided; instead, full statistical distributions (light gray for GTP, dark gray for GDP) are shown overlaid with the average lifetimes. The length of protofilaments was fixed and equal to 6 dimers in all simulations. **(D)** Top view of representative microtubule end structures in both nucleotide states for selected lateral interaction energy values. Gray arrowheads indicate protofilament clusters.

We next calculated the distribution of protofilament cluster lifetimes from the moment they formed to their complete or partial dissociation (Fig. 2C). We observed larger lifetimes and, consequently, broader lifetime distributions for increasing *U*_*lat*_. Interestingly, the size, the number and the average lifetime of clusters were consistently smaller for the GDP-microtubule ends compared to the GTP-microtubule ends, irrespective of the lateral interaction energy (Fig. 2A–C). In our previous MD study,^45^ GDP-protofilaments were observed to be more restricted in terms of radial and tangential bending, explaining why they would, on average, form fewer, smaller and shorter-lived clusters compared to the GTP-protofilaments at a given *U*_*lat*_. Figure 2D visualizes the described quantitative trends by showing representative snapshots of microtubule ends in both nucleotide states.

We also observed that larger protofilament clusters tend to adopt straighter conformations, supporting our hypothesis that protofilament clusters are polymerization intermediates. Moreover, the free energy required to straighten a protofilament cluster decreases rapidly with the cluster size and shows a weaker dependence on the protofilament length (Fig. S2; see next section for the description of the single-cluster setup). This effect is also expected within the “tug-of-war” concept: the microtubule end gains additional energy by forming lateral bonds between neighboring protofilaments; this energy is then spent on “forcing” the protofilament clusters into straighter conformations away from their equilibrium shape.

Altogether, these results demonstrate that a wide spectrum of statistical distributions of protofilament cluster numbers, sizes and lifetimes can be achieved by fine-tuning the spring-like properties of protofilaments and their lateral interaction energies. Moreover, this fine-tuning does not require large-scale conformational changes in the flared structure of microtubule ends, which aligns well with previous^40–42^ and our own cryoET measurements (Fig. 1B and Fig. S1A). It is hence plausible that GTP-microtubule ends that form more, larger and longer-lived protofilament clusters are also more polymerization-prone because their statistical ensemble is more similar to a fully straight lattice.

### Nucleotide state affects protofilament length via asymmetric strain in clusters and protofilament ruptures

To avoid studying finite size effects, we asked if and how the dynamics and the distribution of clusters would change with the length of protofilaments, *L*_*PF*_. To this end, we repeated the above simulations for *L*_*PF*_ between 3 and 9 dimers and calculated 2D parametric diagrams for the average number of clusters (Fig. S3A,B) and the average fraction of protofilaments in clusters (Fig. S3C,D). These simulations showed that, for small to moderate lateral strengths (*U*_*lat*_ ≲ 40 kJ/mol), the propensity to form clusters generally decreased with increasing *L*_*PF*_. This implies that for small *L*_*PF*_, the free energy spent on straightening the protofilaments, and the free energy gained from forming lateral bonds between the protofilaments approximately compensate each other, enabling more frequent cluster formation. At the same time, the straightening energy increases faster with *L*_*PF*_ than the lateral energy decreases, shifting the balance towards more splayed end conformations for large *L*_*PF*_. We believe that there are two reasons for the faster increase of the curved-to-straight free energy barrier: (i) a higher enthalpic contribution due to twist accumulation in longer protofilaments, and (ii) a higher entropic contribution due to a larger phase space available to longer protofilaments.

For high lateral strengths (*U*_*lat*_ ≳ 40 kJ/mol), the propensity to form clusters remained constant or even increased with increasing *L*_*PF*_; however, the simulated microtubule ends were increasingly distorted and unstable. When inspecting simulation trajectories, we observed events in which protofilaments at the edges of clusters would spontaneously rupture and dissociate. Such rupture events were rarely observed during the characteristic lifetime of protofilament clusters (Fig. 2C), but still occurred sufficiently often within the 30 × 200 μs timescale of our simulations. Interestingly, these events mainly occurred on the left side of a cluster when viewed from within the lumen (see the example in Fig. 3A). This unexpected observation prompted us to study the statistics of rupture events in more detail.

**Figure 3.**
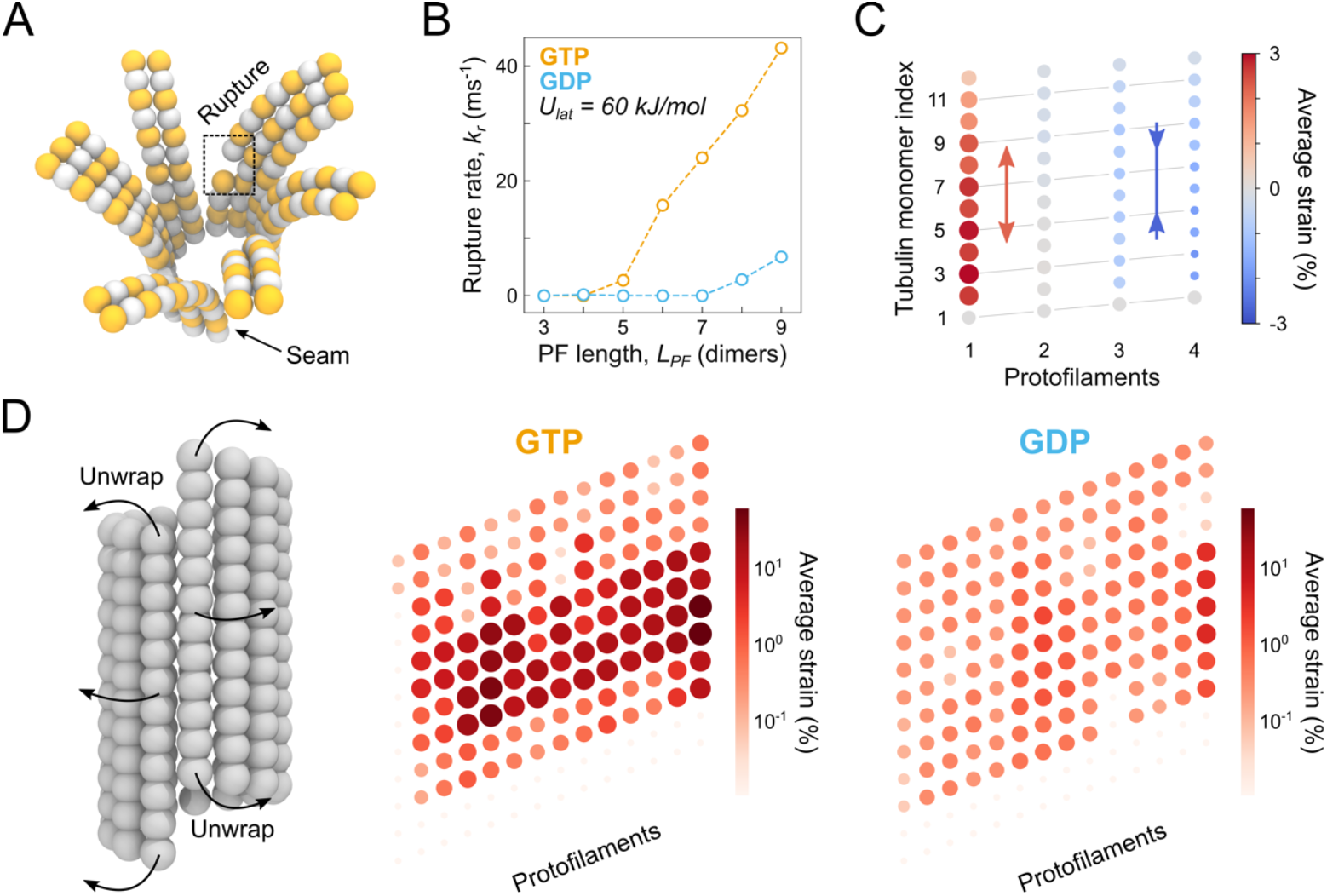
**(A)** Snapshot from one of our CG simulations demonstrating a rupture event (dashed square). **(B)** The rate of rupture events *k*_*r*_ plotted as a function of protofilament length *L*_*PF*_ and nucleotide state (orange for GTP and cyan for GDP). The lateral interaction strength was fixed at *U*_*lat*_ = 60 kJ/mol (see Fig. S4A,B for the full 2D parametric diagrams). **(C)** The average relative strain along GDP-protofilaments in a cluster of size 4 and *L*_*PF*_ = 6 dimers relative to that in the initial straight configuration. Each circle corresponds to a tubulin monomer while its color and size denote the magnitude and the sign of strain, respectively. The lateral and longitudinal bonds were replaced with harmonic potentials to prevent dissociation. **(D)** The average relative strain along protofilaments in a full GTP- and GDP-microtubule lattice of *L*_*PF*_ = 6 dimers relative to that in the initial straight configuration, when unwrapped onto a 2D lattice representation. The lateral interaction strength was fixed at *U*_*lat*_ = 60 kJ/mol. Each circle corresponds to a tubulin monomer while the color and size denote the magnitude of strain. Log-scale was chosen to emphasize the difference in mechanical frustration between GTP- and GDP-microtubule ends.

First, we quantified how frequently ruptures occurred in the GTP- and GDP-microtubule trajectories depending on *L*_*PF*_. While only very few ruptures were seen for both nucleotide states when *U*_*lat*_ ≲ 40 kJ/mol, the rupture rate in GTP-microtubule ends steeply increased when *U*_*lat*_ ≳ 40 kJ/mol and *L*_*PF*_ ≳ 5 dimers (Fig. 3B; see Fig. S4A,B for the full 2D parametric diagrams). When combined with Fig. 2A–C and Fig. S3, these data suggest that cluster formation might be linked to protofilament rupture and subsequent dissociation of tubulin dimers or oligomers via a negative feedback mechanism.

To test this idea, and in particular the reciprocal causal relationship, we constructed GTP-clusters of size between 1 and 4 protofilaments and performed independent simulations of these clusters for different *L*_*PF*_. To keep the clusters intact over the course of the simulation, we made all lateral and longitudinal bonds harmonic. Figure 3C shows the average distribution of longitudinal mechanical strain in a cluster of size 4 protofilaments and *L*_*PF*_ = 6 dimers. This distribution was strongly asymmetric, with the highest stretching strain localized on the left side (where most ruptures had occurred in our simulations) and the highest compression strain localized on the right side of the cluster when viewed from within the lumen. The asymmetry became more pronounced with increasing either *L*_*PF*_ or the cluster size (see Fig. S5 for the full 2D parametric diagrams). Thus, these simulations of “indestructible” clusters demonstrate that excess stretching strain correlates with the location of protofilament rupture, regardless of the cluster size and *L*_*PF*_.

Finally, we quantified how the longitudinal mechanical strain causing protofilament rupture was distributed across the complete microtubule end and how it depended on the nucleotide state. To this end, we re-analyzed the simulation datasets shown in Fig. 2 as follows: each trajectory was truncated just before the first rupture occurred, and the average strain per dimer was calculated over all the independent and truncated trajectories. Unexpectedly, unlike in the single cluster case (Fig. 3C), the average strain in the entire microtubule was predominantly positive, i.e. the protofilaments were, on average, overstretched relative to their initial straight configurations. More specifically, the maximum average strain measured was +58.3% and +4.0% for the GTP- and GDP-microtubule ends, respectively, while the minimum average strain was less than -0.04% in both cases. To visualize the difference between the strain distributions in the GTP- and GDP-microtubule ends more clearly, we neglected the small fraction of negative strains and used a log-scale (Fig. 3D). The GTP-microtubule lattice was, on average, much more mechanically frustrated – despite the known increased radial and tangential softness of its protofilaments.^45^ Moreover, the localization of excess strain within the GTP-microtubule end coincided well with the most frequent location of ruptures, namely near the lattice shaft and distant from the protofilament tips. We believe that this is because (i) protofilaments are softer to stretching deformations than compression ones,^60,61^ and (ii) unlike in Fig. 3C, the lateral and longitudinal bonds are breakable. This observation further indicates that, despite the asymmetric deformation behavior localized in isolated “indestructible” clusters, the short-lived nature of clusters in a more realistic, full microtubule simulation (Fig. 2), combined with protofilament exchange among them, leads to an almost uniform probability distribution of longitudinal bond breakages across the protofilaments, leading to a more even distribution in *L*_*PF*_.

Taken together, these data demonstrate an interesting phenomenon: while GTP-microtubule ends form larger and longer-lived protofilament clusters more frequently (Fig. 2 and Fig. S3), the resulting excess mechanical frustration in the clusters leads to more frequent protofilament ruptures (Fig. 3, Fig. S4 and Fig. S5), thus affecting the shape of the microtubule ends. It is conceivable that the ability to form and maintain sufficiently large protofilament clusters correlates with a reduction of *L*_*PF*_ and vice versa. The fact that this reciprocal relationship is strongly nucleotide-dependent (Fig. S3 and Fig. S4) sets constraints on the potential mechanism of microtubule assembly. In particular, because GTP-microtubule ends favor configurations with large and long-lived clusters, which decreases the average *L*_*PF*_, newly incoming dimers are more likely to form a sufficient number of lateral bonds to get accommodated into the lattice. Conversely, a gradual loss of the GTP cap due to hydrolysis shifts the conformational preference towards smaller and less stable clusters, which reduces the mechanical frustration but increases the average *L*_*PF*_, thereby raising the free energy barrier to convert individual protofilaments into a straight microtubule lattice.

### Electron tomography confirms predicted structural differences at the ends of growing and shortening microtubules

The self-assembly mechanism proposed above enables several predictions that can be tested experimentally. First, the ends of growing microtubules should have shorter protofilaments. Second, they should have more protofilament clusters. Third, the difference in *L*_*PF*_ as well as the difference in the frequency and the size of clusters between growing and shortening microtubule ends should be significant yet subtle, because this would otherwise render dynamic instability energetically unfeasible.

To test these three predictions, we performed cryoET using dynamic microtubules reconstituted from purified tubulin and GMPCPP-stabilized seeds attached to electron microscopy grids. To image growing microtubules, we froze the grids after several minutes of incubation with high tubulin concentration. To image shortening microtubules, we diluted tubulin to below 2 μM and froze the grids after 30–45 seconds. We then performed cryoET and denoised the tomograms using the Cryo-CARE approach.^58^ After determining the polarity of microtubules, we manually segmented plus ends and traced their protofilaments (Fig. 4A, Fig S6; see Materials and Methods). From these 3D traced models, we first obtained the samples of *L*_*PF*_ measured from the first segment bending away from the microtubule cylinder (n=1113 and n=922 for growing and shortening end, respectively). Analysis of these samples revealed a wide distribution that was shifted towards longer protofilaments for shortening microtubule ends (Fig. 4B), consistent with previous reports.^40,41^ A two-sample Kolmogorov-Smirnov test additionally showed that the two samples do not belong to the same unknown distribution (statistic *D*_*K*−*S*_ = 0.17 and p-value *p*_*K*−*S*_ 10^−12^).

**Figure 4.**
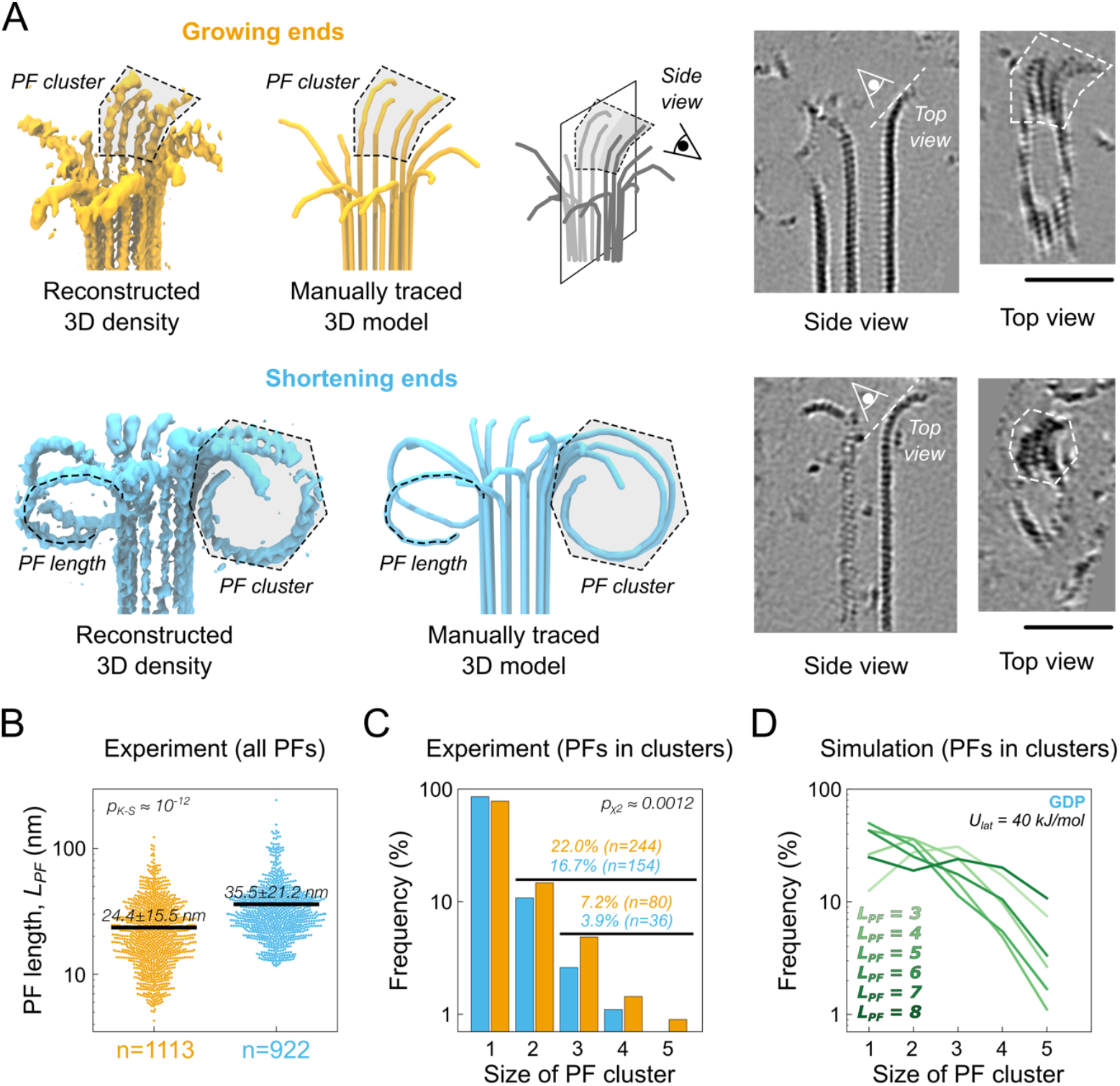
**(A)** Segmented and 3D rendered volumes and manually traced 3D models showing growing (orange, left) and shortening (cyan, right) microtubule ends. A typical protofilament cluster is marked with a light gray dashed area. Also shown are tomographic slices with a cluster in cross-section (side view), and parallel to the initial segment of a cluster flaring out of the microtubule cylinder (top view). Scale bars are 50 nm. **(B)** Distribution of protofilament lengths for cryoET samples imaged in the presence of soluble tubulin under growing (orange) and shortening (cyan) conditions. Shown are raw data points (dots) corresponding to individual protofilaments and mean values (black lines). Log-scale was chosen to visualize the two distinct distributions on a single scale. The value in the upper left corner reports the results of a Kolmogorov-Smirnov test and its p-value. **(C)** Distribution of protofilament cluster sizes including single protofilaments from the experimental datasets shown to the left. The value in the upper right corner reports the results of Pearson’s *χ*^2^ test and its p-value. The values *n* report the numbers of individual protofilaments in a category (all clusters and large clusters of 3 or more protofilaments). **(D)** Same as in **(C)** but calculated from the simulated ensembles of GDP-microtubule ends shown in Fig. 2. Shown are the cluster size statistics for multiple *L*_*PF*_ and *U*_*lat*_ = 40 kJ/mol.

Further, we excluded all protofilaments that had no proximal neighbors based on their normalized mutual overlap (see Materials and Methods and Fig. S7) and interpreted the remaining fraction as protofilament clusters. We sorted all protofilaments according to the size of the clusters they belonged to and constructed a contingency table summarizing the counts for all cluster sizes in both polymerization states. We observed 22.0% (n=244) and 16.7% (n=154) of all protofilaments in clusters at growing and shortening microtubule ends, respectively (Fig. 4C). Additionally, 7.2% (n=80) of protofilaments at growing ends formed large clusters with 3 or more protofilaments, whereas only 3.9% (n=36) of large clusters were observed at shortening ends. Pearson’s *χ*^2^ test confirmed that the difference in the observed counts of protofilaments belonging to clusters of a particular size at growing or shortening microtubule ends is statistically significant (statistic *χ*^2^ = 18.1 and p-value *p* ≈ 0.0012).

We also computed the cluster size statistics from our simulations. Figure 4D shows the distribution of cluster sizes for the GDP-microtubule ends simulated at *U*_*lat*_ = 40 kJ/mol and for different *L*_*PF*_, demonstrating that it is, in principle, possible to select such a pair of *U*_*lat*_ and *L*_*PF*_ for our model to approximately reproduce the experimental values (Fig. 4C, cyan). Despite this favorable agreement, we must note that it is not fully quantitative because, while in our simulations all protofilaments have the same length, in experiments, the protofilament length distribution at the microtubule end is very ragged (Fig 4B). Nevertheless, it is remarkable that the experimentally observed cluster sizes lie well within the range covered by our CG simulations. Finally, whereas the distributions of both *L*_*PF*_ and cluster sizes differ only subtly (Fig. 4B,C), we have now been able to detect this difference and show its statistical significance, thanks to the improved resolution provided by Cryo-CARE. Taken together, our cryoET results are in remarkable agreement with the predictions delivered by the CG simulations.

## Conclusions

Based on our findings presented here, we propose a new mechanism of microtubule assembly, which we term *conformational selection*. It synergizes the previous cryoET experiments by McIntosh, Gudimchuk and others^40–42^ as well as the large-scale atomistic MD simulations of microtubule ends.^44–46^ Figure 5 schematically illustrates its key differences to the previously proposed mechanism, which we term *induced fit* for consistency.

**Figure 5.**
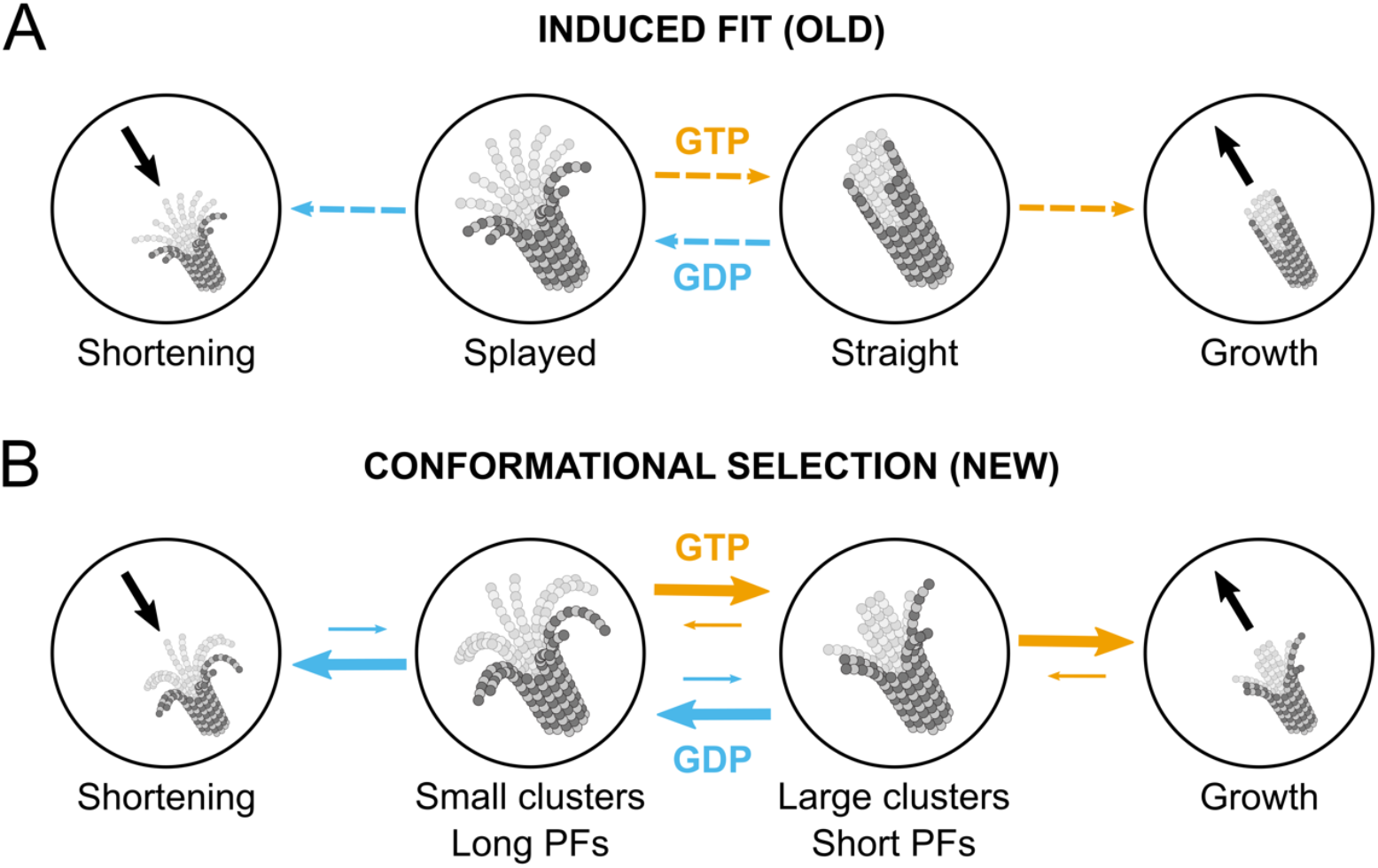
**(A)** Schematic illustration of the induced fit **(A)** and the conformational selection **(B)** mechanisms. Note that the solid arrows in **(B)** denote kinetic rates according to our model, where thicker and longer arrows correspond to higher kinetic rates. In contrast, the dashed arrows in **(A)** simply show the direction of preferred transitions based on the nucleotide state.

The induced fit mechanism (Fig. 5A) postulates that the shape of protofilaments at the microtubule end is directly controlled by the nucleotide state. Upon GTP hydrolysis, the initially straight and growing microtubule end begins to coil inside-out and shorten, producing splayed protofilaments and oligomeric disassembly products in solution. In contrast, the new conformational selection mechanism (Fig. 5B) postulates that, irrespective of the nucleotide state, all microtubule ends are splayed and can form protofilament clusters (Fig. 1B and Fig. S1A) driven by the “tug-of-war” between protofilament elasticity and lateral interactions^44,45^ (Fig. 1A). The nucleotide state modulates the mechanical flexibility of each protofilament, resulting in larger (GTP) or smaller (GDP) clusters (Fig. 2 and Fig. S3). Clusters of long GTP-protofilaments are subject to excess mechanical strain and protofilament ruptures (Fig. 3, Fig. S4 and Fig. S5), leading to GTP-protofilaments being, on average, shorter but forming larger clusters than GDP-protofilaments (Fig. 4). These fast, microsecond “tug-of-war” dynamics, combined with tension-induced, nucleotide-dependent protofilament ruptures at sub-millisecond timescales as well as tubulin binding/unbinding at millisecond timescales (20–40 ms^54,62,63^ at 10 μM tubulin) drive the conformational selection for polymerization-prone configurations showing higher similarity to a straight microtubule end (Fig. 5B, middle). GTP hydrolysis reduces the probability of growth by decreasing the average cluster size, resulting in long, “ram’s horn”-like GDP-protofilaments and pairs thereof, thereby triggering microtubule catastrophe. Besides providing striking agreement with our cryoET data (Fig. 4), this novel model emphasizes and explains the stochastic nature of microtubule self-assembly and catastrophe driven by GTP hydrolysis.

This conformational selection mechanism may help explain microtubule behavior in various physiological and pathological contexts or prompt reassessment studies of fundamental aspects of microtubule dynamics that are now taken for granted. Important examples include the regulation of microtubule dynamics by force and microtubule-associated proteins. For example, tensile forces stabilize the kinetochore-microtubule attachment in mitosis, stall microtubule disassembly and induce microtubule rescue.^64,65^ The kinetochore could bundle and straighten neighboring GDP-protofilaments of the disassembling microtubule end into a more GTP-like state (increased clustering), which could provide a possible explanation for the kinetochore’s ability to remain attached to microtubule ends under force.^66–68^ Further, the conformational selection mechanism could offer a leverage for multivalent microtubule-binding oligomers of the human Ndc80 or budding yeast Dam1 complexes to fine tune the polymerization dynamics by controlling protofilament clusters, without either the microtubule end or the kinetochore having to undergo large conformational changes.

The conformational selection mechanism could also explain why human end-binding proteins EB1/3 and their fission yeast homologues Mal3 accelerate microtubule growth.^69,70^ Since both bind microtubule lattices in between neighboring protofilaments,^48,71^ we speculate that they either directly promote the formation of new protofilament clusters or stabilize existing or potentially emerging protofilament clusters against dissociation. The latter scenario appears more plausible as Mal3 has a higher affinity for a hydrolysis intermediate of GTP-tubulin, with the maximum occupancy site located slightly behind the growing end,^69^ a location to which it recruits other proteins in massive comet-like assemblies.^72^ Within the conformational selection model, binding of EB1/3 and Mal3 would shift the dynamic “tug-of-war” equilibrium towards a straighter microtubule lattice and, thus, accelerate conformational maturation of the growing end.

Finally, binding of taxol has been shown to stabilize microtubules below or near stoichiometric equivalence with tubulin dimers both in vitro^73^ and in vivo;^74^ however, the stabilization mechanism is still debated. More recently, taxol binding has been shown to invert the conformational change that normally occurs in response to GTP hydrolysis, producing expanded and more heterogeneous microtubule lattices in high-resolution cryo-electron microscopy densities obtained by the Nogales lab.^47,75^ We and others have also confirmed in a series of atomistic MD studies that expanded tubulin conformations result in softer protofilaments and microtubule lattices.^45,57,61^ Assuming that these softening effects are hallmarks of a GTP-like state of tubulin, we speculate that taxol binding increases protofilament radial and tangential flexibility, shifting the equilibrium towards larger clusters of shorter protofilaments. Indeed, a recent cryoET study has reported that growing microtubule ends treated with 10 nM of taxol feature even shorter protofilaments than growing microtubule ends in the control experiment,^41^ though protofilament clusters have not been analyzed.

This type of regulation by modulating the stability of self-assembly intermediates (protofilament clusters) might not be unique to taxol. Studies have been performed showing that microtubules disassemble faster in the presence of high concentrations of Mg^2+^, and that adding substantial amounts of Mg^2+^ creates longer protofilament curls^5,76,77^ and increases the work transferred by these curls in optical tweezers assays.^8^ Thus, it cannot be ruled out that certain mutations not interfering with lateral and longitudinal lattice interfaces or the nucleotide binding pocket exploit the conformational selection mechanism to alter microtubule dynamic instability, for example β:T238A causing faster growth in hyperstable yeast and human microtubule phenotypes^78,79^ or β:D417H/β:R262H linked to ocular motility disorder in humans and also causing faster growth.^80^

From the simulation perspective, our CG model (Fig. 1C,D, Fig. S1 and Materials and Methods) accounts for a number of structural and dynamical properties that were previously obtained through accurate atomistic MD simulations of tubulin oligomers and complete microtubule ends.^45^ These simulations set strict physical constraints on the type of degrees of freedom and the nature and strength of tubulin-tubulin interactions. For example, previous CG models did not allow for the possibility of protofilaments deviating from their radial planes and did not consider the bending-torsional coupling within each protofilament, and therefore did not predict intermediate protofilament cluster states. Here, we show that a rigorous, discrete elastic model optimized against high-resolution atomistic simulations can overcome this limitation and reach excellent agreement with experiment. Nevertheless, our CG model still contains a number of approximations. First, our CG mapping (Fig. 1C and Fig. S1B) might miss important degrees of freedom that have not yet been resolved by electron microscopy or have been observed in all-atom MD but not recognized as functionally relevant. Second, in our CG model, all elementary protofilament deformations are harmonic, and only local correlations are considered, neglecting nonlinear mechanical effects or long-range interaction components. Third, although our CG model has been parameterized using extensive atomistic MD trajectories (∼400 μs of cumulative sampling), these do not account for potentially relevant conformational changes occurring at longer timescales. Many of these limitations will be overcome in future by more elaborated CG mappings and parametrization schemes as well as by more exhaustive atomistic MD simulations.

It is noteworthy that the excellent agreement between our simulations and experimental data (Fig. 4C,D) also results from an improved signal-to-noise ratio provided by Cryo-CARE denoising.^58^ This procedure has allowed us to obtain higher resolution in 3D compared to previous studies.^40–42^ However, other limitations imposed by cryoET of flexible protofilaments remain. For example, missing wedge artifacts can result in under-sampling of clustered protofilaments that are located in unfavorable orientations and can only be overcome by dual-axis tomography.^81^ In addition, manual segmentation and tracing of tubulin coordinates may introduce a human bias which is, however, reduced thanks to the increased signal-to-noise ratio. Automated segmentation and tracing of denoised volumes will likely increase the reliability and throughput of our method in future.

Several important questions and concerns remain to be addressed. Although our simulation and cryoET data lead to the unexpected discovery that it is mainly the conformation and stability of protofilament clusters – and not the overall shape – that determine the polymerization state of a microtubule end, we have not yet determined its critical structural ensemble for which the probabilities of growth and shortening are equal. Microtubule “aging” – the increase of the catastrophe probability with time – is another interesting but mechanistically not yet understood phenomenon, and it is currently unclear whether it can be explained within the conformational selection model or, alternatively, is caused by accumulating lattice defects.^81^ Additionally, our CG model so far cannot describe mixed nucleotide lattices which are also difficult to resolve experimentally. We believe that accurate CG models accounting for kinetic transitions caused by GTP hydrolysis and guided by the computational and structural findings presented here will contribute to our understanding of the mechanisms of catastrophe and rescue. To answer these crucial questions, we need to unify high-resolution electron and optical microscopy and biochemistry with advanced computational approaches in an integrative structural biology framework.

## Materials and Methods

### Preparation of in vitro microtubule samples for cryoET

Microtubules were polymerized using purified porcine brain tubulin (Cytoskeleton Inc). First, we prepared stabilized microtubule seeds by polymerizing 25 μM tubulin (with or without addition of 40% digoxigenin (DIG)-labeled tubulin) in the presence of 1 mM GMPCPP, a slowly hydrolyzable GTP analog. After 30 min of polymerization at 37°C, seeds were sedimented in a Beckman Airfuge, resuspended on ice in MRB80 (80 mM K-Pipes pH 6.9 with 4 mM MgCl2 and 1 mM EGTA). After 20 min of depolymerization on ice, seeds were re-polymerized in the presence of freshly added 1 mM GMPCPP, sedimented again, resuspended in MRB80 with 10% glycerol at room temperature, aliquoted and snap-frozen in liquid nitrogen.

Samples with growing microtubules ends were made by polymerizing microtubules in the presence of seeds, 15 μM tubulin, 1 mM GTP, and 5 nm gold nanoparticles at 37°C for 30 min in a dry bath. 3.5 μL of this mixture were added to a freshly glow-discharged lacey carbon grid (Agar Scientific) suspended in the chamber of a Leica EM GP2 plunge-freezer equilibrated at 37°C and 99% relative humidity. After the addition of microtubules, the grid was blotted from the back side and immediately frozen in liquid ethane.

Samples with shortening microtubules were made with DIG-labeled seeds attached to the surface of either 1.2/1.3 Quantifoil grids or silanized holey silicon oxide grids (SPI Supplies) as described previously.^72^ Briefly, the grids were incubated with anti-DIG IgG, washed with MRB80, then incubated with DIG-labeled GMPCPP-stabilized microtubule seeds and suspended in the chamber of a Leica EM GP2 plunge-freezer equilibrated at 37°C and 99% relative humidity. 3 μL of 20 μM tubulin in MRB80 supplemented with 1mM GTP were added to the grid and incubated in the chamber for 7 min. After that, 30 μL of pre-warmed MRB80 were added to the grid to induce microtubule depolymerization. Dilution buffer was supplemented with 5 nm gold nanoparticles before addition to the grid. Majority of the buffer dripped off from the grid, leaving 3-4 μL that were blotted off from the back side during 30–45 s after the addition of the buffer. Immediately after the blotting, the grid was frozen in liquid ethane. All grids were stored in liquid nitrogen until further use.

### Data acquisition

Growing microtubules (three separate samples) were imaged using a JEM-3200FSC electron microscope (JEOL) equipped with a K2 Summit direct electron detector (Gatan) and an in-column energy filter operated in zero-loss imaging mode with a 30-eV slit width. Images were recorded at 300 kV with a nominal magnification of 10,000x, resulting in the pixel size of 3.668 Å at the specimen level. Automated image acquisition was performed using SerialEM software.^82^ A subset of growing microtubules (one additional sample) was imaged using a Titan Krios microscope equipped with a Gatan K3 electron detector at the Netherlands Center for Electron Nanoscopy (NeCEN, Leiden, Netherlands), at 300 kV with a nominal magnification of 26000x, resulting in the pixel size of 3.28 Å at the specimen level. Automated image acquisition was performed using SerialEM. Shortening microtubules (two separate samples) were imaged using a Titan Krios microscope (FEI) equipped with a Gatan K2 electron detector (NeCEN). Automated image acquisition was performed using Tomography software (Thermo Fisher). Images were recorded at 300 kV with a nominal magnification of 33,000x, resulting in the pixel size of 4.24 Å at the specimen level. Energy filtering for all data collected at NeCEN was performed at post-processing. In all cases, we recorded bi-directional tilt series ranging from 0° to ±60° with a tilt increment of 2°, the total electron dose of 100 e−/Å2 and the target defocus set to -4 μm.

### Image processing of microtubule ends

Tomograms were reconstructed and denoised as described previously,^58,72^ using tomograms generated with even and odd frames after alignment with MotionCor2,^83^ and tilt series alignment and back projection performed in IMOD.^84^

Further analysis was limited to microtubule plus-ends. Microtubule polarity was determined using visual inspection of moiré patterns of protofilaments after Fourier filtering.^85^ Since microtubules were polymerized from GMPCPP seeds, most of them contained 14 protofilaments, making this analysis unambiguous in the majority of cases. The polarity was confirmed by observing microtubule cross-sections after the denoising procedure and noting the direction of protofilament “wedges”.^86^ Each protofilament at a plus-end was manually traced using 3dmod,^84^ while observing the accuracy of the segmentation by simultaneously visualizing the rendered denoised experimental density and the model in the *isosurface* view, as described previously.^87^

### Structural analysis of protofilament clusters in cryoET reconstructions

To filter out and analyze protofilament clusters, we calculated the overlap between the volumes occupied by neighboring protofilaments. We equidistantly distributed spheres along the protofilament traces with an increment of 0.1 nm. We set the equilibrium distance between two protofilaments in a cluster and the sphere radius to 5.34 nm and 3.20 nm, respectively. We further observed that manual tracing errors introduced distortions in the protofilament traces, leading to unnaturally high local curvature. This was likely due to the low resolution of tomograms. To mitigate the impact of these errors, we introduced weights *w*_*i*_ linearly increasing from 0 to 1 along each protofilament trace’s length. These weighting factors did not affect cluster detection in “good” cases but increased the probability to detect clusters with high distortion. The rationale behind this weighting is as follows: If the ends of two distorted protofilaments are close together, they are very likely to be in a cluster owing to the high out-of-plane protofilament stiffness. Finally, the total volume overlap between two neighboring protofilament traces, Ω, was calculated as the sum of the weighted sphere volume overlaps Δ*V*_*i*_ normalized by the sum of the weighted maximum volume overlaps Δ*V*_*i,max*_ (assuming ideally straight protofilaments):

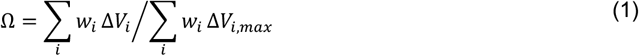

where Δ*V*_*i*_ ≤ Δ*V*_*i,max*_. The threshold for the volume overlap in a protofilament cluster was chosen to be 10% (see Fig. S7). This value indicates that at least 50% of the weighted linear distances between protofilaments deviate by less than 20% from the ideal case.

### Discrete elastic rod model of the microtubule lattice

We used a discrete elastic rod (DER) representation to model individual protofilaments.^88^ Originally devised for modeling macroscopic elastic materials and creating visual effects in animated film industries, the DER formalism produces strikingly realistic dynamics of hair, trees and viscous fluids.^89–91^ This discrete differential geometry approach is designed to handle arbitrary deformable configurations, diverse cross-sections and dynamic complexities. Here, we applied it to model the microscopic microtubule end structure as a set of 14 coupled DERs (Fig. 1C).

Each configuration of a DER was represented by a centerline consisting of *N* + 1 nodes ***r***^***i***^, where 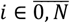, connected by *N* edges. Each edge *j*, where 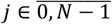, was associated with a material frame 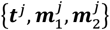 that formed a right-handed orthonormal triad, with the tangential vector ***t***^*j*^ being oriented along the edge. The material frame described the orientation of the rod and, together with the twist-free (Bishop) frame,^89^ was used to define the DER twist. Stretching, bending and twisting deformations along the DER were represented by (i) the edge vector lengths *e*^*j*^ = ‖***e***^*j*^ ‖ = ‖***r***^*j*^ − ***r***^*j*−1^ ‖, (ii) the discrete integrated curvature vectors (*k****b***)^*i*^, where *k*^*i*^ is the discrete integrated curvature and ***b***^***i***^ is the discrete binormal vector and (iii) the discrete integrated twists *m*^*i*^ = *θ*^*i*^ − *θ*^*i*−1^, where *θ*^*i*^ is the angle between the material and the Bishop frame. To calculate this twist, we used the reference frame as described previously.^90^ Notably, in the DER formalism, stretching and compression are the properties of edges (pairs of nodes), while bending and twisting are the properties of nodes (pairs of edges), except for the terminal nodes that cannot be assigned a curvature or twist. In addition, twisting and bending of only neighboring edges relative to one another are correlated, i.e. no long-range effects along the protofilament are assumed.

The potential function *U* describing the energetics of the microtubule end was composed of the elastic energy of protofilaments and the lateral and longitudinal interaction energies between neighboring protofilaments (Fig. 1D). Following the canonical DER approach, the elastic energy was further composed of the stretching *U*_*S*_, twisting *U*_*t*_ and bending *U*_*b*_ energies. An additional potential *U*_*tb*_ was introduced to explicitly account for the positive linear correlation between bending and twisting as this correlations was observed in our previous atomistic simulations of tubulin dimers and protofilaments.^45,59^ All of these potentials were harmonic with respect to the elementary deformations, and their full mathematical expressions are given in Table S1.

By definition, forces in the DER formalism act only on nodes and edge twist angles (i.e. the canonical coordinates), whereas a dimer in our system was described by 3 consecutive nodes in the DER (Fig. 1C). Consequently, the lateral interaction between two dimers in neighboring protofilaments was modeled as a 3-node interaction, with each pair of interacting nodes contributing one third. To this end, 2 virtual sites were introduced for every node in the protofilament located at distances −*r* and +*r* on the axis defined by 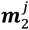 of the associated edge (Fig. 1D), and the interaction between the virtual sites was modeled using a Morse potential. To account for volume exclusion, an additional repulsive potential between laterally neighboring nodes was introduced that was modeled using the repulsive part of a Lennard-Jones potential. Finally, we modeled the longitudinal interaction between two consecutive tubulin dimers in a protofilament by replacing the harmonic potential describing the stretching/compression of the edge corresponding to α-tubulin with another Morse potential (Fig. 1D). The full mathematical expressions for the lateral *U*_*lat*._ and longitudinal *U*_*long*_ interaction energies are given in Table S1.

### Brownian Dynamics simulations

To quantify the time evolution of the microtubule end, we integrated the overdamped Langevin equations of motion for the node positions ***r***^*i*^ and the edge twist angles *θ*^*j*^ in every protofilament *k* simultaneously:

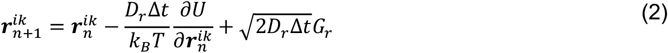

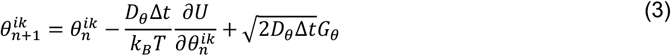

where ∆*t* is the integration time between steps *n* + 1 and *n, U* = *U*_*S*_ + *U*_*t*_ + *U*_*b*_ + *U*_*tb*_ + *U*_*lat*_ + *U*_*long*_ is the full potential function, *D*_*r*_ and *D*_*θ*_ are the translational and rotational diffusion constants for the nodes and twist angles, *k*_*B*_ is the Boltzmann constant, *T* is the temperature, and *G*_*r*_ and *G*_*θ*_ are Gaussian distributed random numbers with zero mean and unit variance. *D*_*r*_∆*t* and *D*_*θ*_∆*t* were adjusted such as to roughly reproduce the relaxation timescales of both the single protofilaments and the entire microtubule from our previous simulations (*D*_*r*_∆*t* ≈ 2 × 10^−4^ nm^2^ and *D*_*θ*_∆*t* ≈ 2 × 10^−4^ rad^2^).^45^

### Mapping and parametrization of the coarse-grained microtubule end model

There are two distinct stages to constructing a CG model: mapping and parametrization. The mapping procedure defines the resolution of a CG model and how well it captures the properties of structure, mechanics and symmetry.^92^ In our case, the task was to find a mapping of the DER’s centerline to the atomistic structure of a protofilament. To decouple stretching/compression deformations from bending deformations, this centerline should pass through groups of atoms in the protofilament structure such that the mutual distances between these groups do not change during protofilament bending, i.e. through “hinges” connecting tubulin monomers around which dimers twist and bend. Early structural studies identified helices H8 and H11’ as key interaction sites between the monomers in a tubulin dimer and the dimers in a protofilament.^16,17^ It was later shown computationally that most protofilament bending and twisting was enabled by these small and robust “hinges”.^31,45,57^

We used the residues βH8:249–264 and αH11’:405–411 and αH8:251–266 and βH11’:395–401 to define the nodes ***r***^*ik*^ corresponding to intra- and inter-dimer interfaces in the DER model, respectively. We further specified the material frames 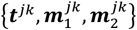 for the edges to complete the mapping. To this end, we drew an imaginary line that connected each edge’s center of mass (COM) with the main microtubule axis and that was orthogonal to that edge. We then selected a group of atoms within a sphere of radius 0.25 nm located 2 nm away from the edge’s COM on this line. We finally constructed a monomer vector between the edge’s COM and this group of atoms for every tubulin monomer in the system. Finally, the material vectors 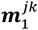 and ***t***^*jk*^ were defined as the normalized monomer and edge vectors, respectively. The material vectors 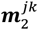 were naturally defined to form a right-handed triad with 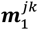 and ***t***^*jk*^.

Methods for the parametrization stage are diverse, and the models they produce can have distinct levels of accuracy.^92^ In our case, the optimization task for a single protofilament was to obtain CG parameters that, given the above mapping, best reproduce the global essential dynamics of bending-torsional fluctuations of protofilaments.^45^ To this end, we used fuzzy self-tuning particle swarm optimization (FST-PSO)^93,94^ to minimize a custom-made objective function employing (i) Earth mover’s distance (EMD)^95^ as a metric to compare all-atom MD and CG distributions of local deformations and (ii) Principal Component Analysis (PCA)^96,97^ as a metric to compare all-atom MD and CG distributions of the global essential dynamics (see Fig. S1C). The objective function was as follows:

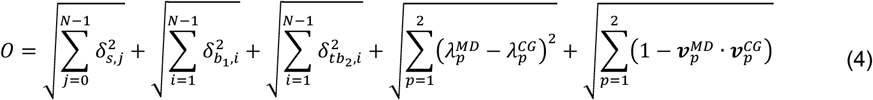

where *δ*_*s,j*_ is the EMD between the all-atom MD and CG distribution of the *j*-th edge vector length, 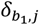 is the EMD between the all-atom MD and CG distribution of the first principal curvature for the *i*-th non-terminal node, 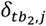 is the EMD between the all-atom MD and CG joint 2D distribution of the second principal curvature and twist for the *i*-th non-terminal node and *λ*_*p*_ and ***v***_*p*_ are the eigenvalues and the eigenvectors of the first 2 PCA components describing the protofilament dynamics in all-atom MD or CG representation.^45^ The first and second principal curvatures are projections of the discrete integrated curvature vectors (*k****b***)^*i*^ onto the material frames associated with the *i*-th node (see Jawed et al.^88^ for the exact definitions) and describe radial and tangential bending of the DER, respectively.

There were a total of 18 parameters describing the deformation of a PF: 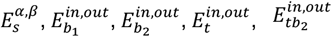 (stretching, bending, twisting and twist-bending moduli, respectively) and 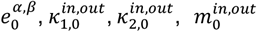 (equilibrium edge lengths, curvatures and twists, respectively), where the indices *α* and *β* denote edges corresponding to α- and β-tubulin and the indices *in* and *out* denote nodes corresponding to intra- and inter-dimer interfaces, respectively. Since the bending, twisting and twist-bending coupling terms build a positive definite quadratic form, their rigidity parameters must additionally satisfy the condition 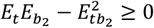 for every node. To simplify the problem, 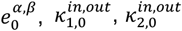 and 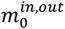 were not optimized explicitly; instead, they were set to the mean values obtained from the all-atom MD simulations converted to the DER representation. To achieve fast convergence and avoid sampling local minima of the 10-dimensional rigidity parameter space, we set up and ran 2200 independent FST-PSO optimizations, each starting from a random vector. The search space was limited to a physically reasonable range of [10^2^, 4 × 10^4^] for each parameter (regardless of the units), and the optimization was terminated when the objective function did not decrease for 1000 consecutive steps. For each of the optimized parameters, we collected a distribution of the 2200 FST-PSO solutions and calculated the mean value, which was then chosen to be the final optimization result (see Fig. S1D and Table S2). We finally obtained 2 separate sets of the optimized parameters for both GTP and GDP systems to account for the effect of the nucleotide state.

As specified above, the lateral and longitudinal dimer-dimer interaction energies were approximated with Morse potentials, with a repulsive Lennard-Jones component added to the lateral energy. The Morse potentials contained a total of 6 parameters: the interaction energies 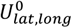, the scaling terms *a*_*lat,long*_ and the equilibrium distances 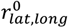, where the indices *lat* and *long* denote lateral and longitudinal interaction interfaces, respectively. To simplify the problem, the repulsive Lennard-Jones constant *σ* was fixed and set to 5.34 nm, the COM-COM distance between neighboring dimers in a microtubule lattice. The equilibrium distances 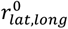 were not optimized explicitly; instead, they were fixed and set to the values obtained from the all-atom free energy calculations. The optimization was performed only for parameters 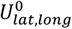 and *a*_*lat,long*_ as described above for the DER using a simplified objective function:

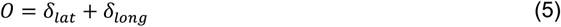

where *δ*_*lat*._ and *δ*_*long*_ are the EMDs between the all-atom MD and CG distributions of the COM-COM distance for a pair of laterally and longitudinally coupled dimers or monomers, respectively (see Table S2). Importantly, we later used the lateral interaction energy parameter 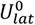 as an external free parameter in our study to manually control the “tug-of-war” balance (Fig. 1A), and a dimensionless scaling factor was introduced into the system’s potential function to control its magnitude.

To optimize the lateral parameters, we used the free energy calculations from our previous study (see Fig. S1E),^45^ while we performed new free energy calculations for the longitudinal parameters (see Fig. S1F). We finally obtained 4 separate sets of the optimized parameters for the lateral interaction for both GTP and GDP systems and both homotypic and seam interfaces as well as 2 separate parameter sets for the longitudinal interaction for both GTP and GDP systems. In total, the atomistic datasets used to parametrize our CG model of the microtubule end comprised approximately 400 μs of cumulative sampling.

### All-atom free energy calculations for the longitudinal dimer-dimer bond

GTP- and GDP-tubulin oligomer systems composed of two longitudinally coupled dimers were prepared, solvated, neutralized with 150 mM KCl and equilibrated as described previously.^45^ We employed the umbrella sampling approach^98^ in conjunction with the weighted histogram analysis method (WHAM).^99,100^ We first defined the COM-COM distance between α- and β-tubulin belonging to the inter-dimer interface as the reaction coordinate. The biasing potential was tuned to be 4000 kJ/mol/nm^2^. To cover the full range of inter-dimer interactions, the COM-COM distance range between 4.2 and 6.5 nm was split into windows, each being separated by 0.05 nm. This partitioning of the reaction coordinate space yielded sufficient overlap between neighboring windows in the absolute majority of cases. The equilibration run was used for seeding the umbrella simulations, where each seed was separated from all the others by at least 50 ns in time. Seeding structures for those windows that were not initially covered by the equilibration simulation of the oligomer systems were derived from neighboring windows located 0.05 nm away in the reaction coordinate space. Each window was then simulated for 500 ns. We calculated the free energy profiles and their uncertainties using WHAM and Bayesian bootstrapping of the complete histograms scaled by inefficiency factors.^100^

### Data analysis and availability

All MD simulations were done using GROMACS 2023.^101^ All post-processing calculations and data analyses were done with GROMACS internal tools, Python 3.9^102^, Numpy v1.26^103^ and SciPy v1.11.^104^ Graphs were produced using Matplotlib v3.8.2^105^ and Seaborn v0.13.^106^ All structure and cryoET density manipulations were performed using Chimera v1.17^107^ or Visual Molecular Dynamics (VMD) v1.9.3.^108^ The VMD software was further used for visualization of microtubule end structures. All CG simulations of microtubule ends and protofilament clusters were performed using a custom Python code accelerated with Numba.^109^ CryoET data presented in Fig. 4A were deposited to EMDB (EMD-52025 for growing microtubule ends, EMD-52026 shortening microtubule ends).

## Acknowledgments

We thank Wiel Evers (Delft University of Technology) and Christoph Diebolder (Netherlands Centre for Electron Nanoscopy – NeCEN) for their excellent technical assistance in data acquisition for cryoET. In addition, M.K., M.I. and H.G. acknowledge the Max Planck Society as well as the Deutsche Forschungsgemeinschaft (DFG, German Research Foundation) – Project-ID 449750155 – RTG 2756, Project A1 for supporting this research. V.A.V. acknowledges the start-up funding provided by the Queen Mary University of London (SBC8VOL2). M.I. acknowledges the start-up funding provided by the School of Life Sciences, University of Dundee.

## Notes

### Competing Interest Statement

The authors have declared no competing interest.

### Summary of Updates

additional supplementary figures and movies; improved figures and statistical analysis; minor text improvements

## References

1. Baas, P. W. Microtubules and neuronal polarity: lessons from mitosis. Neuron 22, 23–31 (1999).

2. Roll-Mecak, A. The tubulin code in microtubule dynamics and information encoding. Dev. Cell 54, 7–20 (2020).

3. Gudimchuk, N. B. & McIntosh, J. R. Regulation of microtubule dynamics, mechanics and function through the growing tip. Nat. Rev. Mol. Cell Biol. 22, 777–795 (2021).

4. Dogterom, M. & Yurke, B. Measurement of the force-velocity relation for growing microtubules. Science 278, 856–860 (1997).

5. Grishchuk, E. L., Molodtsov, M. I., Ataullakhanov, F. I. & McIntosh, J. R. Force production by disassembling microtubules. Nature 438, 384–388 (2005).

6. Volkov, V. A. et al. Long tethers provide high-force coupling of the Dam1 ring to shortening microtubules. Proc. Natl. Acad. Sci. U. S. A. 110, 7708–7713 (2013).

7. Driver, J. W., Geyer, E. A., Bailey, M. E., Rice, L. M. & Asbury, C. L. Direct measurement of conformational strain energy in protofilaments curling outward from disassembling microtubule tips. Elife 6, e28433 (2017).

8. Murray, L. E., Kim, H., Rice, L. M. & Asbury, C. L. Working strokes produced by curling protofilaments at disassembling microtubule tips can be biochemically tuned and vary with species. Elife 11, e83225 (2022).

9. Mitchison, T. & Kirschner, M. Dynamic instability of microtubule growth. Nature 312, 237–242 (1984).

10. Caplow, M. & Shanks, J. Mechanism of the microtubule GTPase reaction. J. Biol. Chem. 265, 8935–8941 (1990).

11. Caplow, M., Ruhlen, R. L. & Shanks, J. The free energy for hydrolysis of a microtubule-bound nucleotide triphosphate is near zero: all of the free energy for hydrolysis is stored in the microtubule lattice. J. Cell Biol. 127, 779–788 (1994).

12. Drechsel, D. N. & Kirschner, M. W. The minimum GTP cap required to stabilize microtubules. Curr. Biol. 4, 1053–1061 (1994).

13. Roostalu, J. et al. The speed of GTP hydrolysis determines GTP cap size and controls microtubule stability. Elife 9, e51992 (2020).

14. Estévez-Gallego, J. et al. Structural model for differential cap maturation at growing microtubule ends. Elife 9, e50155 (2020).

15. LaFrance, B. J. et al. Structural transitions in the GTP cap visualized by cryo-electron microscopy of catalytically inactive microtubules. Proc. Natl. Acad. Sci. U. S. A. 119, e2114994119 (2022).

16. Nogales, E., Wolf, S. G. & Downing, K. H. Structure of the alpha beta tubulin dimer by electron crystallography. Nature 391, 199–203 (1998).

17. Löwe, J., Li, H., Downing, K. H. & Nogales, E. Refined structure of αβ-tubulin at 3.5 Å resolution 1 1Edited by I. A. Wilson. J. Mol. Biol. 313, 1045–1057 (2001).

18. Simon, J. R. & Salmon, E. D. The structure of microtubule ends during the elongation and shortening phases of dynamic instability examined by negative-stain electron microscopy. J. Cell Sci. 96 (Pt 4), 571–582 (1990).

19. Mandelkow, E. M., Mandelkow, E. & Milligan, R. A. Microtubule dynamics and microtubule caps: a time-resolved cryo-electron microscopy study. J. Cell Biol. 114, 977–991 (1991).

20. Chrétien, D., Fuller, S. D. & Karsenti, E. Structure of growing microtubule ends: two-dimensional sheets close into tubes at variable rates. J. Cell Biol. 129, 1311–1328 (1995).

21. Müller-Reichert, T., Chrétien, D., Severin, F. & Hyman, A. A. Structural changes at microtubule ends accompanying GTP hydrolysis: information from a slowly hydrolyzable analogue of GTP, guanylyl (alpha,beta)methylenediphosphonate. Proc. Natl. Acad. Sci. U. S. A. 95, 3661–3666 (1998).

22. Manuel Andreu, J., Garcia de Ancos, J., Starling, D., Hodgkinson, J. L. & Bordas, J. A synchrotron X-ray scattering characterization of purified tubulin and of its expansion induced by mild detergent binding. Biochemistry 28, 4036–4040 (1989).

23. Buey, R. M., Díaz, J. F. & Andreu, J. M. The nucleotide switch of tubulin and microtubule assembly: a polymerization-driven structural change. Biochemistry 45, 5933–5938 (2006).

24. Rice, L. M., Montabana, E. A. & Agard, D. A. The lattice as allosteric effector: structural studies of alphabeta- and gamma-tubulin clarify the role of GTP in microtubule assembly. Proc. Natl. Acad. Sci. U. S. A. 105, 5378–5383 (2008).

25. Gigant, B. et al. Structural basis for the regulation of tubulin by vinblastine. Nature 435, 519– 522 (2005).

26. Nawrotek, A., Knossow, M. & Gigant, B. The determinants that govern microtubule assembly from the atomic structure of GTP-tubulin. J. Mol. Biol. 412, 35–42 (2011).

27. Ayaz, P., Ye, X., Huddleston, P., Brautigam, C. A. & Rice, L. M. A TOG:αβ-tubulin complex structure reveals conformation-based mechanisms for a microtubule polymerase. Science 337, 857–860 (2012).

28. Rakshit, D., Daily, K. M. & Blume, D. Natural and unnatural parity states of small trapped equal-mass two-component Fermi gases at unitarity and fourth-order virial coefficient. arXiv [cond-mat.quant-gas] 587–590 (2011) doi:10.1126/science.1230582.

29. Gebremichael, Y.Chu, J.-W. & Voth, G. A. Intrinsic bending and structural rearrangement of tubulin dimer: molecular dynamics simulations and coarse-grained analysis. Biophys. J. 95, 2487–2499 (2008).

30. Bennett, M. J. et al. Structural mass spectrometry of the alpha beta-tubulin dimer supports a revised model of microtubule assembly. Biochemistry 48, 4858–4870 (2009).

31. Grafmüller, A. & Voth, G. A. Intrinsic bending of microtubule protofilaments. Structure 19, 409– 417 (2011).

32. André, J. R. et al. The state of the guanosine nucleotide allosterically affects the interfaces of tubulin in protofilament. J. Comput. Aided Mol. Des. 26, 397–407 (2012).

33. VanBuren, V., Cassimeris, L. & Odde, D. J. Mechanochemical model of microtubule structure and self-assembly kinetics. Biophys. J. 89, 2911–2926 (2005).

34. Molodtsov, M. I. et al. A molecular-mechanical model of the microtubule. Biophys. J. 88, 3167– 3179 (2005).

35. Molodtsov, M. I., Grishchuk, E. L., Efremov, A. K., McIntosh, J. R. & Ataullakhanov, F. I. Force production by depolymerizing microtubules: a theoretical study. Proc. Natl. Acad. Sci. U. S. A. 102, 4353–4358 (2005).

36. Efremov, A., Grishchuk, E. L., McIntosh, J. R. & Ataullakhanov, F. I. In search of an optimal ring to couple microtubule depolymerization to processive chromosome motions. Proc. Natl. Acad. Sci. U. S. A. 104, 19017–19022 (2007).

37. Zakharov, P. et al. Molecular and mechanical causes of microtubule catastrophe and aging. Biophys. J. 109, 2574–2591 (2015).

38. Bollinger, J. A., Center for Integrated Nanotechnologies, Sandia National Laboratories, Albuquerque NM 87185, USA. msteve@sandia. gov & Stevens, M. J. Catastrophic depolymerization of microtubules driven by subunit shape change. Soft Matter 14, 1748–1752 (2018).

39. Michaels, T. C., Feng, S., Liang, H. & Mahadevan, L. Mechanics and kinetics of dynamic instability. Elife 9, e54077 (2020).

40. McIntosh, J. R. et al. Microtubules grow by the addition of bent guanosine triphosphate tubulin to the tips of curved protofilaments. J. Cell Biol. 217, 2691–2708 (2018).

41. Gudimchuk, N. B. et al. Mechanisms of microtubule dynamics and force generation examined with computational modeling and electron cryotomography. Nat. Commun. 11, 3765 (2020).

42. Atherton, J., Stouffer, M., Francis, F. & Moores, C. A. Microtubule architecture in vitro and in cells revealed by cryo-electron tomography. Acta Crystallogr. D Struct. Biol. 74, 572–584 (2018).

43. Alexandrova, V. V. et al. Theory of tip structure-dependent microtubule catastrophes and damage-induced microtubule rescues. Proc. Natl. Acad. Sci. U. S. A. 119, e2208294119 (2022).

44. Tong, D. & Voth, G. A. Microtubule simulations provide insight into the molecular mechanism underlying dynamic instability. Biophys. J. 118, 2938–2951 (2020).

45. Igaev, M. & Grubmüller, H. Bending-torsional elasticity and energetics of the plus-end microtubule tip. Proc. Natl. Acad. Sci. U. S. A. 119, e2115516119 (2022).

46. Wu, J. et al. Data-driven equation-free dynamics applied to many-protein complexes: The microtubule tip relaxation. bioRxiv 2024.10.10.617682 (2024) doi:10.1101/2024.10.10.617682.

47. Alushin, G. M. et al. High-resolution microtubule structures reveal the structural transitions in αβ-tubulin upon GTP hydrolysis. Cell 157, 1117–1129 (2014).

48. Zhang, R., Alushin, G. M., Brown, A. & Nogales, E. Mechanistic origin of microtubule dynamic instability and its modulation by EB proteins. Cell 162, 849–859 (2015).

49. Vemu, A. et al. Structure and dynamics of single-isoform recombinant neuronal human tubulin. J. Biol. Chem. 291, 12907–12915 (2016).

50. Zhang, R., LaFrance, B. & Nogales, E. Separating the effects of nucleotide and EB binding on microtubule structure. Proc. Natl. Acad. Sci. U. S. A. 115, E6191–E6200 (2018).

51. Manka, S. W. & Moores, C. A. The role of tubulin-tubulin lattice contacts in the mechanism of microtubule dynamic instability. Nat. Struct. Mol. Biol. 25, 607–615 (2018).

52. van den Berg, C. M. et al. CSPP1 stabilizes growing microtubule ends and damaged lattices from the luminal side. J. Cell Biol. 222, e202208062 (2023).

53. Walker, R. A. et al. Dynamic instability of individual microtubules analyzed by video light microscopy: rate constants and transition frequencies. J. Cell Biol. 107, 1437–1448 (1988).

54. Gardner, M. K. et al. Rapid microtubule self-assembly kinetics. Cell 146, 582–592 (2011).

55. Gardner, M. K., Zanic, M., Gell, C., Bormuth, V. & Howard, J. Depolymerizing kinesins Kip3 and MCAK shape cellular microtubule architecture by differential control of catastrophe. Cell 147, 1092–1103 (2011).

56. Mickolajczyk, K. J., Geyer, E. A., Kim, T., Rice, L. M. & Hancock, W. O. Direct observation of individual tubulin dimers binding to growing microtubules. Proc. Natl. Acad. Sci. U. S. A. 116, 7314–7322 (2019).

57. Fedorov, V. A. et al. Mechanical properties of tubulin intra- and inter-dimer interfaces and their implications for microtubule dynamic instability. PLoS Comput. Biol. 15, e1007327 (2019).

58. Buchholz, T.-O., Jordan, M., Pigino, G. & Jug, F. Cryo-CARE: Content-aware image restoration for cryo-transmission electron microscopy data. in 2019 IEEE 16th International Symposium on Biomedical Imaging (ISBI 2019) 502–506 (IEEE, 2019).

59. Igaev, M. & Grubmüller, H. Microtubule assembly governed by tubulin allosteric gain in flexibility and lattice induced fit. Elife 7, (2018).

60. Wells, D. B. & Aksimentiev, A. Mechanical properties of a complete microtubule revealed through molecular dynamics simulation. Biophys. J. 99, 629–637 (2010).

61. Igaev, M. & Grubmüller, H. Microtubule instability driven by longitudinal and lateral strain propagation. PLoS Comput. Biol. 16, e1008132 (2020).

62. Gudimchuk, N. & Roll-Mecak, A. Watching microtubules grow one tubulin at a time. Proceedings of the National Academy of Sciences of the United States of America vol. 116 7163–7165 (2019).

63. Strothman, C. et al. Microtubule minus-end stability is dictated by the tubulin off-rate. J. Cell Biol. 218, 2841–2853 (2019).

64. Akiyoshi, B. et al. Tension directly stabilizes reconstituted kinetochore-microtubule attachments. Nature 468, 576–579 (2010).

65. Volkov, V. A., Huis In ‘t Veld, P. J., Dogterom, M. & Musacchio, A. Multivalency of NDC80 in the outer kinetochore is essential to track shortening microtubules and generate forces. Elife 7, (2018).

66. Gutierrez, A. et al. Cdk1 phosphorylation of the Dam1 complex strengthens kinetochore-microtubule attachments. Curr. Biol. 30, 4491-4499.e5 (2020).

67. Haase, M. A. B. et al. DASH/Dam1 complex mutants stabilize ploidy in histone-humanized yeast by weakening kinetochore-microtubule attachments. EMBO J. 42, e112600 (2023).

68. Polley, S. et al. Stable kinetochore-microtubule attachment requires loop-dependent Ndc80-Ndc80 binding. EMBO J. 42, e112504 (2023).

69. Maurer, S. P., Fourniol, F. J., Bohner, G., Moores, C. A. & Surrey, T. EBs recognize a nucleotide-dependent structural cap at growing microtubule ends. Cell 149, 371–382 (2012).

70. Maurer, S. P. et al. EB1 accelerates two conformational transitions important for microtubule maturation and dynamics. Curr. Biol. 24, 372–384 (2014).

71. Maurer, S. P., Bieling, P., Cope, J., Hoenger, A. & Surrey, T. GTPgammaS microtubules mimic the growing microtubule end structure recognized by end-binding proteins (EBs). Proc. Natl. Acad. Sci. U. S. A. 108, 3988–3993 (2011).

72. Maan, R. et al. Multivalent interactions facilitate motor-dependent protein accumulation at growing microtubule plus-ends. Nat. Cell Biol. 25, 68–78 (2023).

73. Schiff, P. B., Fant, J. & Horwitz, S. B. Promotion of microtubule assembly in vitro by taxol. Nature 277, 665–667 (1979).

74. Schiff, P. B. & Horwitz, S. B. Taxol stabilizes microtubules in mouse fibroblast cells. Proc. Natl. Acad. Sci. U. S. A. 77, 1561–1565 (1980).

75. Kellogg, E. H. et al. Insights into the distinct mechanisms of action of taxane and non-taxane microtubule stabilizers from cryo-EM structures. J. Mol. Biol. 429, 633–646 (2017).

76. Tran, P. T., Joshi, P. & Salmon, E. D. How tubulin subunits are lost from the shortening ends of microtubules. J. Struct. Biol. 118, 107–118 (1997).

77. O’Brien, E. T., Salmon, E. D. & Erickson, H. P. How calcium causes microtubule depolymerization. Cell Motil. Cytoskeleton 36, 125–135 (1997).

78. Geyer, E. A. et al. A mutation uncouples the tubulin conformational and GTPase cycles, revealing allosteric control of microtubule dynamics. Elife 4, e10113 (2015).

79. Ye, X., Kim, T., Geyer, E. A. & Rice, L. M. Insights into allosteric control of microtubule dynamics from a buried β-tubulin mutation that causes faster growth and slower shrinkage. Protein Sci. 29, 1429–1439 (2020).

80. Ti, S.-C. et al. Mutations in human tubulin proximal to the kinesin-binding site alter dynamic instability at microtubule plus- and minus-ends. Dev. Cell 37, 72–84 (2016).

81. Guyomar, C. et al. Changes in seam number and location induce holes within microtubules assembled from porcine brain tubulin and in Xenopus egg cytoplasmic extracts. Elife 11, (2022).

82. Mastronarde, D. N. SerialEM: A program for automated tilt series acquisition on Tecnai microscopes using prediction of specimen position. Microsc. Microanal. 9, 1182–1183 (2003).

83. Zheng, S. Q. et al. MotionCor2: anisotropic correction of beam-induced motion for improved cryo-electron microscopy. Nat. Methods 14, 331–332 (2017).

84. Kremer, J. R., Mastronarde, D. N. & McIntosh, J. R. Computer visualization of three-dimensional image data using IMOD. J. Struct. Biol. 116, 71–76 (1996).

85. Chrétien, D., Kenney, J. M., Fuller, S. D. & Wade, R. H. Determination of microtubule polarity by cryo-electron microscopy. Structure 4, 1031–1040 (1996).

86. Foster, H. E., Ventura Santos, C. & Carter, A. P. A cryo-ET survey of microtubules and intracellular compartments in mammalian axons. J. Cell Biol. 221, (2022).

87. Ogunmolu, F. E. et al. Microtubule plus-end regulation by centriolar cap proteins. bioRxiv https://doi.org/10.1101/2021.12.29.474442 (2021) doi:10.1101/2021.12.29.474442.

88. Jawed, M. K., Novelia, A. & O’Reilly, O. M. A Primer on the Kinematics of Discrete Elastic Rods. (Springer International Publishing, Cham, Switzerland, 2018).

89. Bergou, M., Wardetzky, M., Robinson, S., Audoly, B. & Grinspun, E. Discrete elastic rods. ACM Trans. Graph. 27, 1–12 (2008).

90. Bergou, M., Audoly, B., Vouga, E., Wardetzky, M. & Grinspun, E. Discrete viscous threads. ACM Trans. Graph. 29, 1–10 (2010).

91. Jawed, M. K. & Reis, P. M. Dynamics of a flexible helical filament rotating in a viscous fluid near a rigid boundary. Physical Review Fluids 2, (2017).

92. Jin, J., Pak, A. J., Durumeric, A. E. P., Loose, T. D. & Voth, G. A. Bottom-up coarse-graining: Principles and perspectives. J. Chem. Theory Comput. 18, 5759–5791 (2022).

93. Kennedy, J. & Eberhart, R. Particle swarm optimization. in Proceedings of ICNN’95 - International Conference on Neural Networks vol. 4 1942–1948 vol.4 (IEEE, 2002).

94. Nobile, M. S. et al. Fuzzy Self-Tuning PSO: A settings-free algorithm for global optimization. Swarm Evol. Comput. 39, 70–85 (2018).

95. Rubner, Y. & Tomasi, C. The earth mover’s distance. in Perceptual Metrics for Image Database Navigation 13–28 (Springer US, Boston, MA, 2001).

96. Amadei, A., Linssen, A. B. & Berendsen, H. J. Essential dynamics of proteins. Proteins 17, 412–425 (1993).

97. de Groot, B. L., van Aalten, D. M., Amadei, A. & Berendsen, H. J. The consistency of large concerted motions in proteins in molecular dynamics simulations. Biophys. J. 71, 1707–1713 (1996).

98. Torrie, G. M. & Valleau, J. P. Nonphysical sampling distributions in Monte Carlo free-energy estimation: Umbrella sampling. J. Comput. Phys. 23, 187–199 (1977).

99. Kumar, S., Rosenberg, J. M., Bouzida, D., Swendsen, R. H. & Kollman, P. A. THE weighted histogram analysis method for free-energy calculations on biomolecules. I. The method. J. Comput. Chem. 13, 1011–1021 (1992).

100. Hub, J. S., de Groot, B. L. & van der Spoel, D. G_wham—A free weighted histogram analysis implementation including robust error and autocorrelation estimates. J. Chem. Theory Comput. 6, 3713–3720 (2010).

101. Abraham, M. et al. GROMACS 2023.2 Manual. Preprint at 10.5281/ZENODO.8134388 (2023).

102. Van Rossum, G. & Drake, F.L., Jr. Python 3 Reference Manual: (Python Documentation Manual Part 2). (Createspace, Scotts Valley, CA, 2009).

103. Harris, C. R. et al. Array programming with NumPy. Nature 585, 357–362 (2020).

104. Virtanen, P. et al. SciPy 1.0: fundamental algorithms for scientific computing in Python. Nat. Methods 17, 261–272 (2020).

105. Hunter, J. D. Matplotlib: A 2D Graphics Environment. Comput. Sci. Eng. 9, 90–95 (2007).

106. Waskom, M. seaborn: statistical data visualization. J. Open Source Softw. 6, 3021 (2021).

107. Pettersen, E. F. et al. UCSF Chimera--a visualization system for exploratory research and analysis. J. Comput. Chem. 25, 1605–1612 (2004).

108. Humphrey, W., Dalke, A. & Schulten, K. VMD: visual molecular dynamics. J. Mol. Graph. 14, 33–8, 27–8 (1996).

109. Lam, S., Pitrou, A. & Seibert, S. Numba: a LLVM-based Python JIT compiler. v7:1-7:6 (2015).

