## Supplementary Information for "Microtubule dynamics are defined by conformations and stability of clustered protofilaments"

**This PDF file includes:**

Figures S1 to S7 with captions
Tables S1 to S2
Legends for Movies S1 to S2
SI References

**Other supporting materials for this manuscript include the following:**

Movies S1 to S2

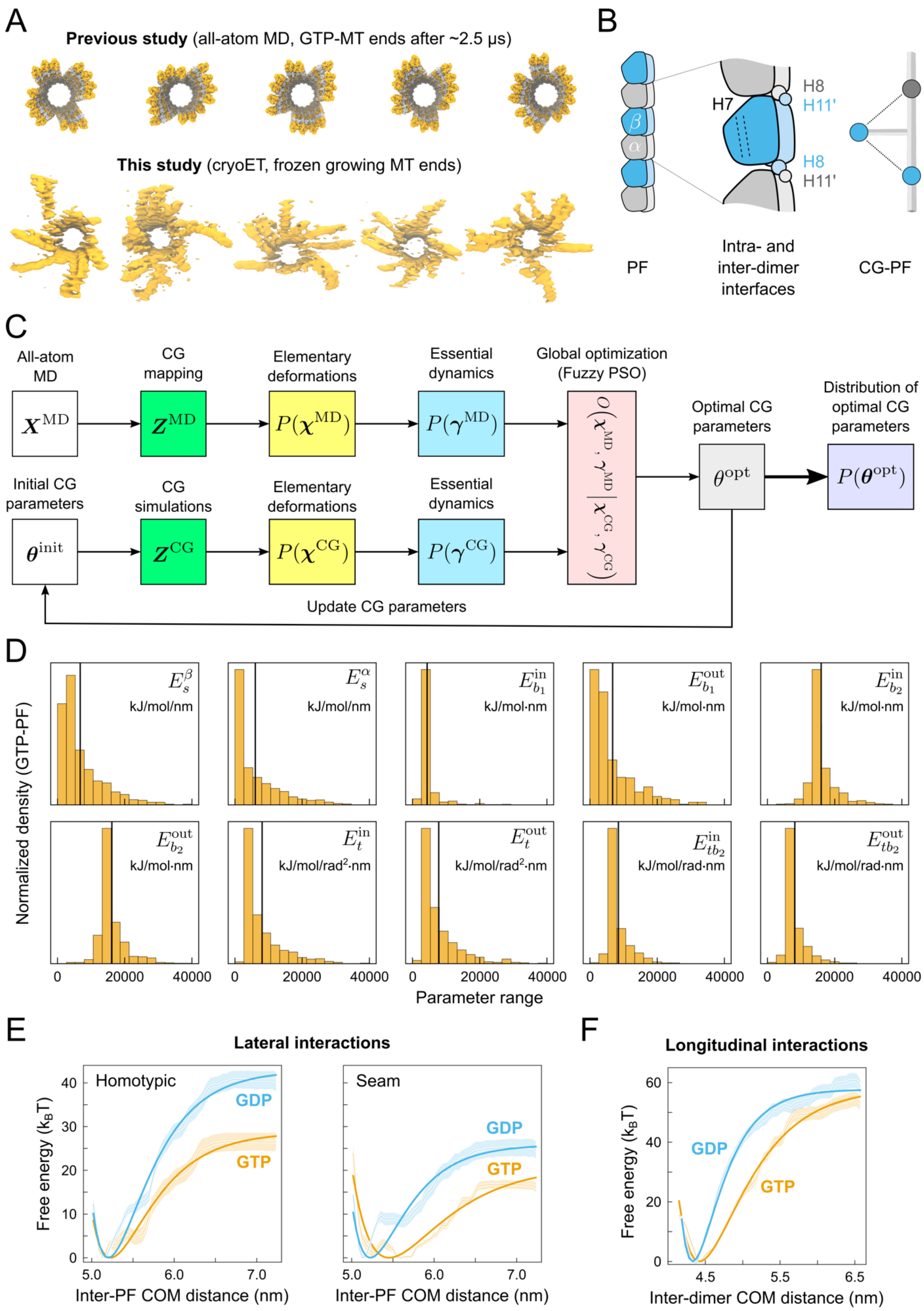
A
Previous study (all-atom MD, GTP-MT ends after ~2.5 μs)
This study (cryoET, frozen growing MT ends)
B
H7
H8
H11'
H8
H11'
PF
Intra- and inter-dimer interfaces
CG-PF
C
All-atom MD
CG mapping
Elementary deformations
Essential dynamics
Global optimization (Fuzzy PSO)
Optimal CG parameters
Distribution of optimal CG parameters
Initial CG parameters
CG simulations
Elementary deformations
Essential dynamics
Update CG parameters
D
Normalized density (GTP-PF)
kJ/mol/nm
kJ/mol/nm
kJ/mol·nm
kJ/mol·nm
kJ/mol·nm
kJ/mol·nm
kJ/mol/rad²·nm
kJ/mol/rad²·nm
kJ/mol/rad·nm
kJ/mol/rad·nm
0
20000
40000
Parameter range
E
Lateral interactions
Homotypic
Seam
GDP
GTP
Free energy (kBT)
Inter-PF COM distance (nm)
F
Longitudinal interactions
GDP
GTP
Inter-dimer COM distance (nm)

**Figure S1. (A)** Comparison of the simulated GTP-microtubule ends from our previous study[1] (top) with exemplary 3D rendered volumes of growing microtubule plus-ends obtained in this study (bottom). **(B)** Schematic illustration of the mapping defining the protofilament centerline. Helices H8 and H11' were used to define the DER nodes, while a group of atoms located within a sphere of radius 2 nm around helix H7 were used to define the material frame. **(C)** Flowchart diagram illustrating the optimization algorithm. **(D)** Distributions of parameters for the GTP-protofilament collected from 2200 independent FST-PSO optimizations (see Table S2 for the full list of parameters for both nucleotide states). The mean values are indicated with black vertical lines. **(E)** Free energy profiles of homotypic (left) and seam-like (right) tubulin–tubulin lateral interactions between two straight, infinitely long protofilaments as a function of the inter-protofilament COM distance.[1] Color coding for GTP and GDP as in Fig. 2. Solid lines indicate the Morse potentials in our CG model optimized against these atomistic data (see Table S2 for the full list of parameters for both nucleotide states). **(F)** Free energy profiles of tubulin-tubulin longitudinal interactions in a short protofilament consisting of two dimers. Color coding as in Fig. 2. Solid lines indicate the Morse potentials in our CG model optimized against these atomistic data (see Table S2 for the full list of parameters for both nucleotide states).

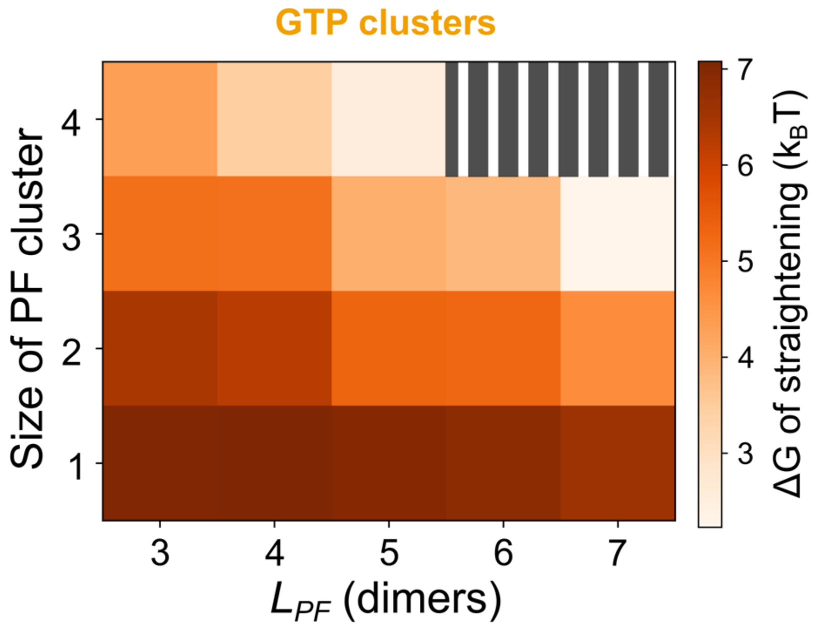


**Figure S2.** Free energy required to straighten a GTP-cluster of size between 1 to 4 protofilaments and $L_{PF}$ between 3 to 7 dimers. The shaded area indicates the parameters for which the formation of clusters was not possible due to excess strain and protofilament rupture. The same single-cluster setup was used as shown in Fig. S5.

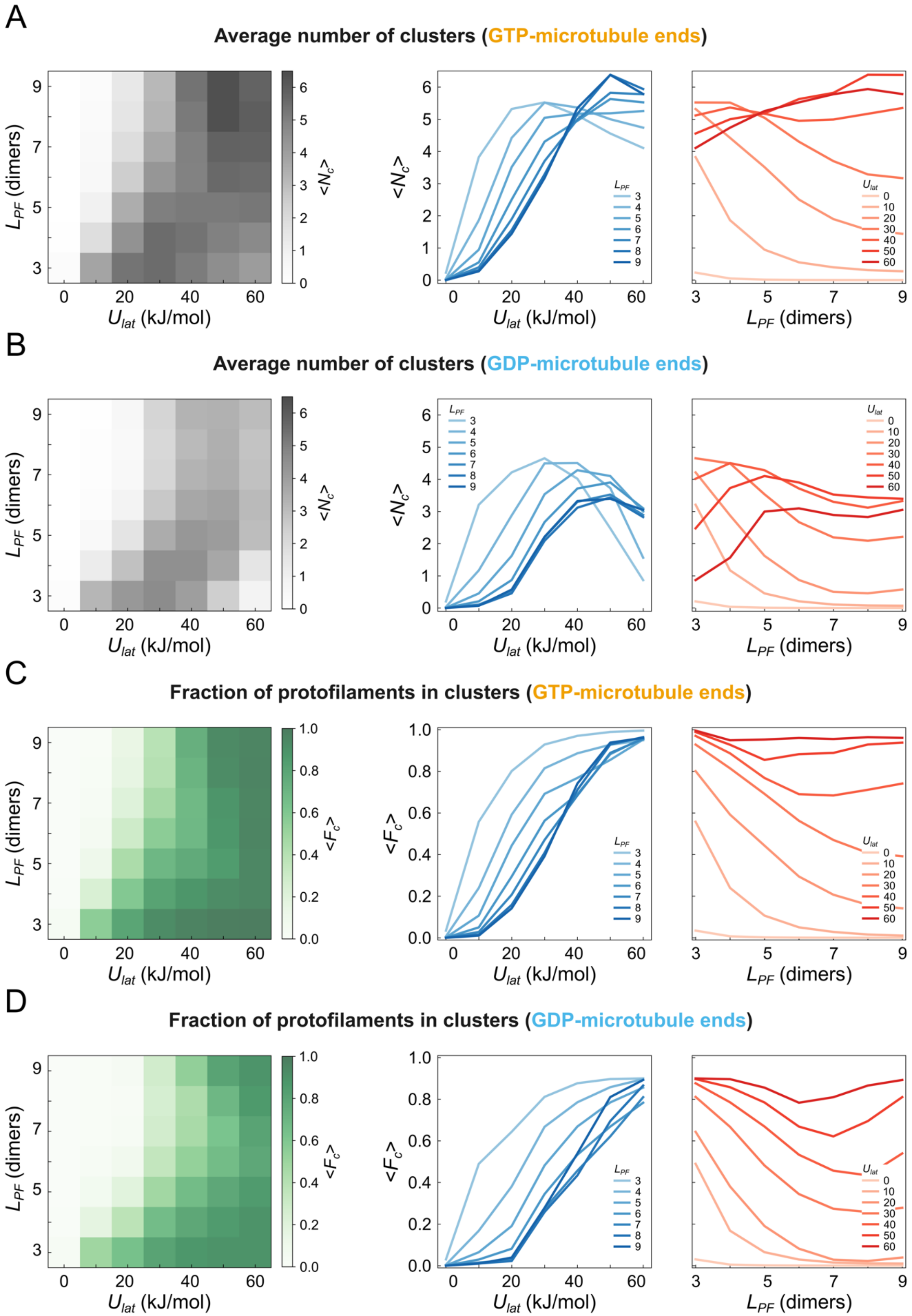
A
Average number of clusters (GTP-microtubule ends)
B
Average number of clusters (GDP-microtubule ends)
C
Fraction of protofilaments in clusters (GTP-microtubule ends)
D
Fraction of protofilaments in clusters (GDP-microtubule ends)
$L_{PF}$ (dimers)
$U_{lat}$ (kJ/mol)
$<N_c>$
$<F_c>$
$L_{PF}$
$U_{lat}$

**Figure S3.** Two-dimensional parametric diagrams of the average number of clusters (<$N_c$>) at **(A)** GTP- and **(B)** GDP-microtubule ends as well as the average fraction of protofilaments (<$F_c$>) in **(C)** GTP- and **(D)** GDP-clusters. Graphs on the left of each panel additionally show the same data as projections on either $U_{lat}$ or $L_{PF}$.

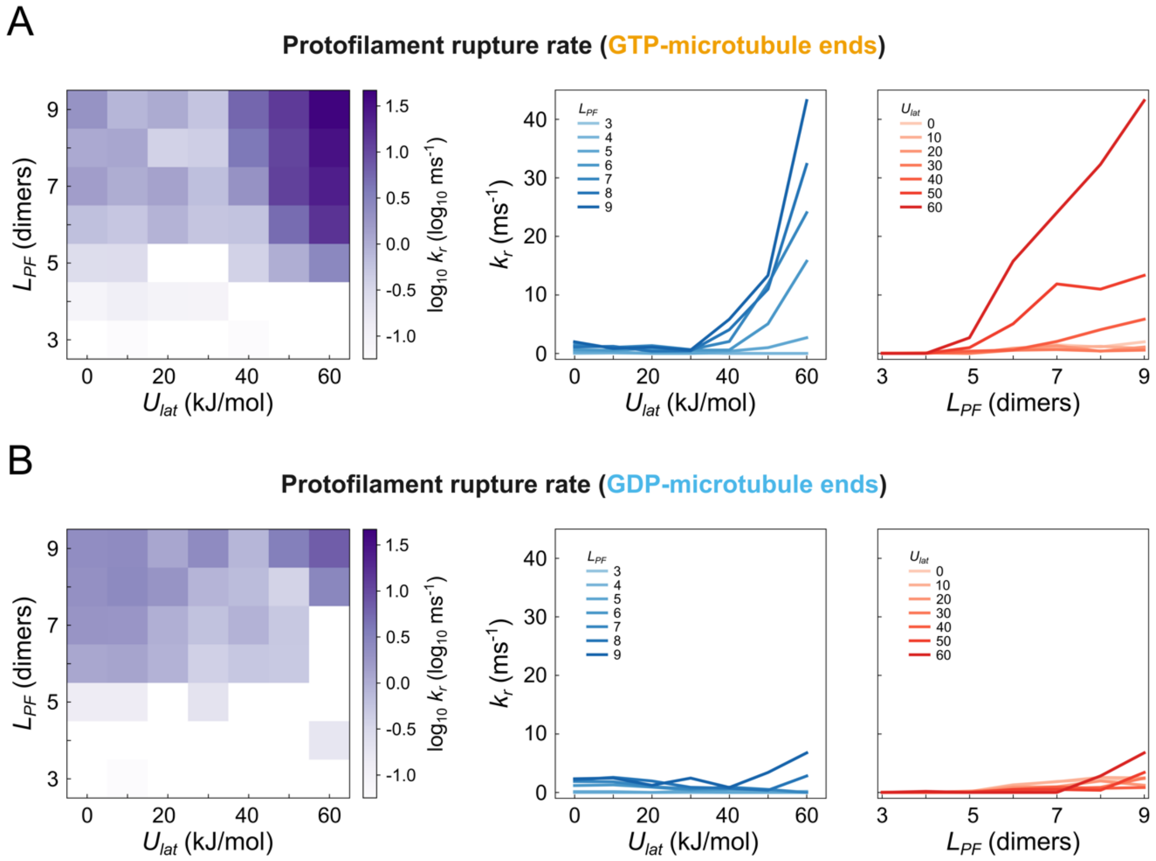


**Figure S4.** Two-dimensional parametric diagrams of the rupture rate ($k_r$) for **(A)** GTP- and **(B)** GDP-protofilaments as a function of lateral interaction strength ($U_{lat}$) and protofilament length ($L_{PF}$). Graphs on the left of each panel additionally show the same data as projections on either $U_{lat}$ or $L_{PF}$. Note that the diagrams use a log-scale for clarity, while the additional graphs use a linear scale to plot the rupture rate.

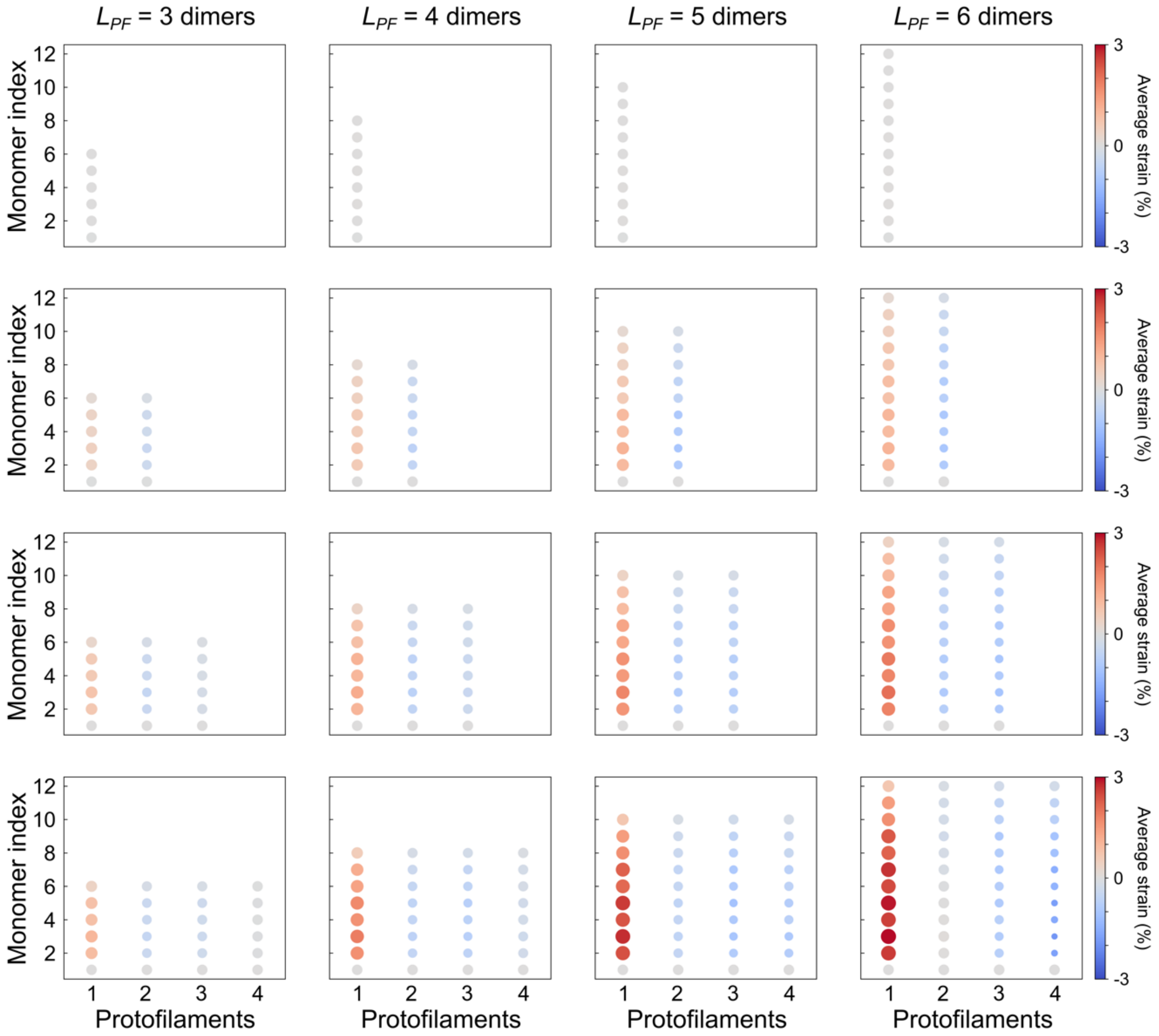


**Figure S5.** Two-dimensional parametric diagrams of the average relative strain along GTP-protofilaments in clusters of size 1 to 4 protofilaments (rows) and length 3 to 6 dimers (columns). Each circle corresponds to a tubulin monomer while its color and size denote the magnitude and the sign of strain, respectively. The lateral and longitudinal bonds were replaced with harmonic potentials to prevent dissociation. The same single-cluster setup was used as shown in Fig. S2 except that here, the lateral and longitudinal bonds were unbreakable.

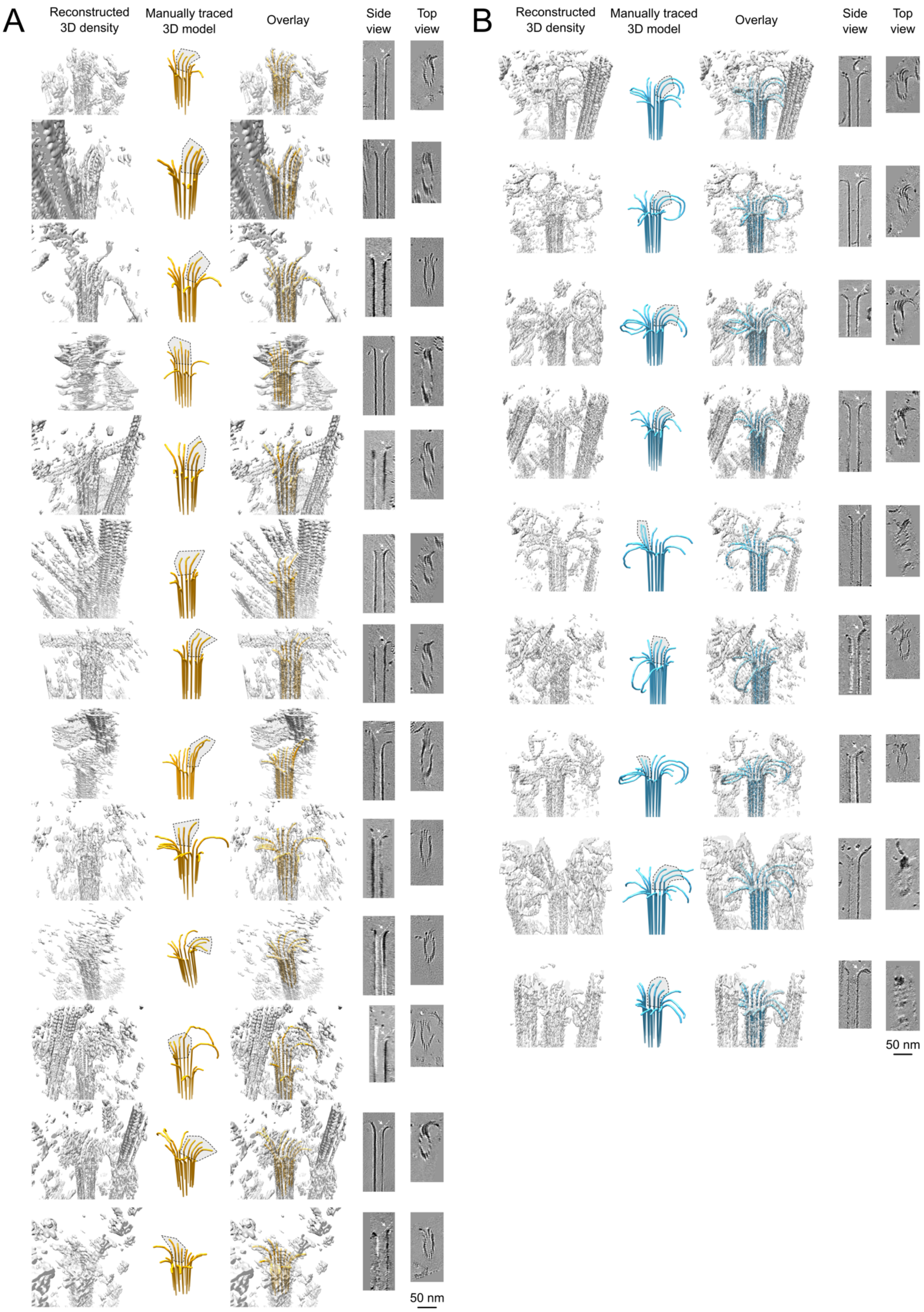

A
Reconstructed 3D density
Manually traced 3D model
Overlay
Side view
Top view
50 nm
B
Reconstructed 3D density
Manually traced 3D model
Overlay
Side view
Top view
50 nm

**Figure S6.** Examples of representative growing **(A)** and shortening **(B)** microtubule ends, in orientations similar to Fig 4A. Shown are from left to right the reconstructed 3D volumes, the manually traced 3D models, the overlay and the side and top views in the corresponding 2D tomograms. Gray areas highlight protofilament clusters. Scale bars are 50 nm.

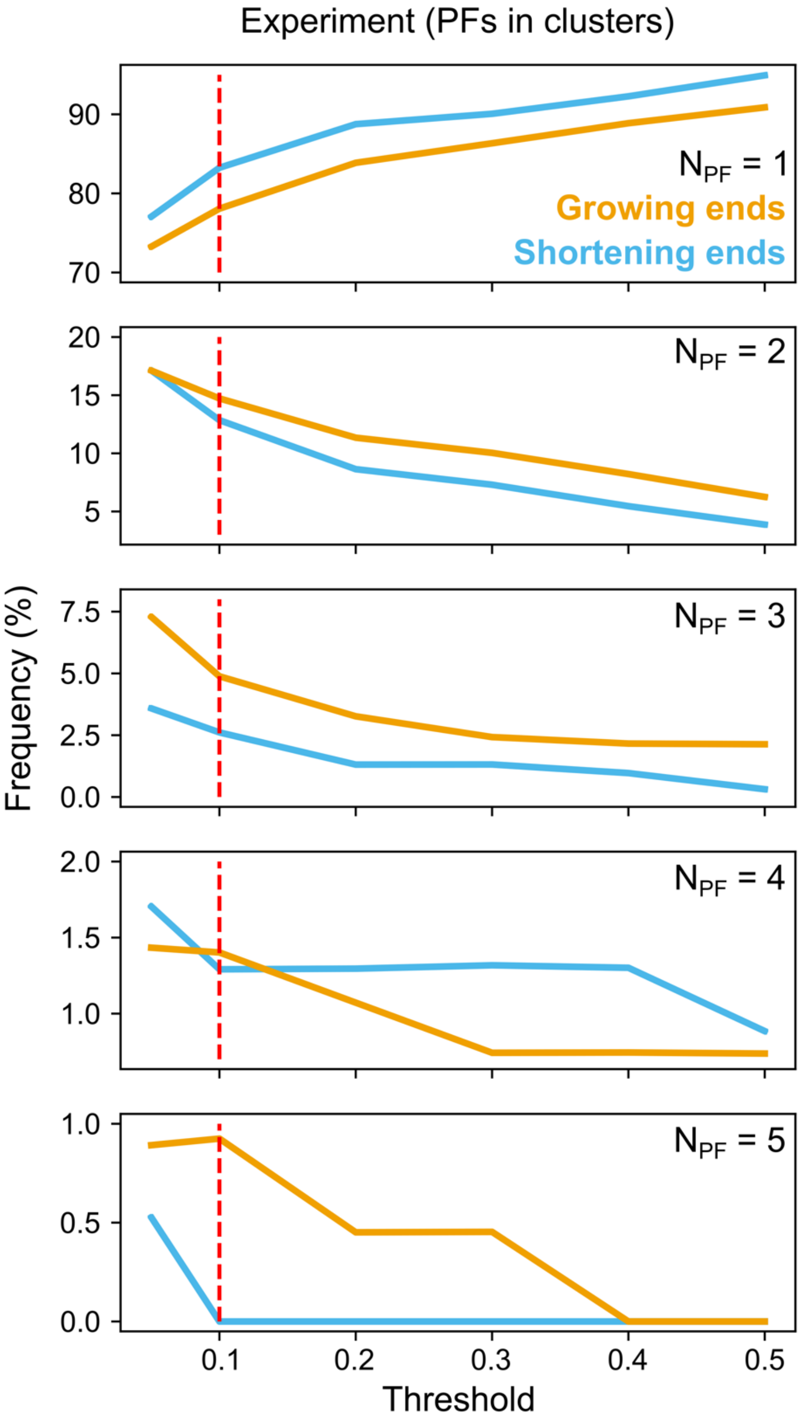


**Figure S7.** The relationship between the threshold in the overlap between neighboring protofilaments, used for defining a protofilament cluster, and the observed frequencies of clusters of various size in our dataset. The red line marks the chosen threshold of 10% used consistently throughout our study. Regardless of the chosen threshold, the quantitative difference between the detected clusters at growing and shrinking ends is preserved.

**Table S1.** Mathematical expressions for the elastic, lateral and longitudinal potential functions defined in Fig. 1D.[2] Also shown are expressions for the auxiliary quantities, i.e. relative extensions $\boldsymbol{\varepsilon}^{\boldsymbol{j}}$, twists $\boldsymbol{m}^{\boldsymbol{i}}$, principal curvatures $\boldsymbol{k}_{\mathbf{1}}^{\boldsymbol{i}}$ and $\boldsymbol{k}_{\mathbf{2}}^{\boldsymbol{i}}$. There are a total of 24 model parameters, from which only 14 parameters were optimized.

| **Potential** | **Expression** | **Schematic** | **Auxiliary quantities** | **Model parameters** |
|---|---|---|---|---|
| *Stretching* | $U_s = \sum_{j=0}^{N-1} E_s^j \lVert\mathbf{e}_0^j\rVert {\varepsilon^j}^2$ | 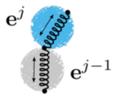 | $\varepsilon^j = \frac{\lVert\mathbf{e}^j\rVert - \lVert\mathbf{e}_0^j\rVert}{\lVert\mathbf{e}_0^j\rVert}$<br>$E_s^j = E_s^\alpha$, if $j$ is even<br>$E_s^j = E_s^\beta$, if $j$ is odd<br>$\lVert\mathbf{e}_0^j\rVert = \lVert\mathbf{e}_0^\alpha\rVert$, if $j$ is even<br>$\lVert\mathbf{e}_0^j\rVert = \lVert\mathbf{e}_0^\beta\rVert$, if $j$ is odd | $E_s^\alpha$ $E_s^\beta$<br>$\lVert\mathbf{e}_0^\alpha\rVert$ $\lVert\mathbf{e}_0^\beta\rVert$ |
| *Twisting* | $U_t = \sum_{i=1}^{N-1} \frac{E_t^i}{l_0^i}\left[m^i - m_0^i\right]^2$ | 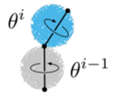 | $m^i = \theta^i - \theta^{i-1}$<br>$l_0^i = \frac{1}{2}\left[\lVert\mathbf{e}_0^{i-1}\rVert + \lVert\mathbf{e}_0^i\rVert\right]$<br>$E_t^i = E_t^{in}$, if $i$ is odd<br>$E_t^i = E_t^{out}$, if $i$ is even<br>$m_0^i = m_0^{in}$, if $i$ is odd<br>$m_0^i = m_0^{out}$, if $i$ is even | $E_t^{in}$ $E_t^{out}$<br>$m_0^{in}$ $m_0^{out}$ |
| *Bending* | $U_b = \sum_{i=1}^{N-1} \frac{E_{b_1}^i}{l^i}\left[\kappa_1^i - \kappa_{1,0}^i\right]^2 + \frac{E_{b_2}^i}{l^i}\left[\kappa_2^i - \kappa_{2,0}^i\right]^2$ | 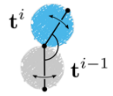 | $\kappa_1^i = \frac{1}{2}\left[\mathbf{m}_2^{i-1} + \mathbf{m}_2^i\right]\cdot(\kappa\mathbf{b})^i$<br>$\kappa_2^i = -\frac{1}{2}\left[\mathbf{m}_1^{i-1} + \mathbf{m}_1^i\right]\cdot(\kappa\mathbf{b})^i$<br>$\mathbf{b}^i = \frac{\mathbf{t}^{i-1}\times\mathbf{t}^i}{\lVert\mathbf{t}^{i-1}\times\mathbf{t}^i\rVert}$<br>$\kappa^i = \mathrm{tg}\left(\frac{\mathbf{t}^{i-1}\wedge\mathbf{t}^i}{2}\right)$<br>$E_{b_{1,2}}^i = E_{b_{1,2}}^{in}$, if $i$ is odd<br>$E_{b_{1,2}}^i = E_{b_{1,2}}^{out}$, if $i$ is even<br>$\kappa_{1,2,0}^i = \kappa_{1,2,0}^{in}$, if $i$ is odd<br>$\kappa_{1,2,0}^i = \kappa_{1,2,0}^{out}$, if $i$ is even | $E_{b_{1,2}}^{in}$ $E_{b_{1,2}}^{out}$<br>$\kappa_{1,2,0}^{in}$ $\kappa_{1,2,0}^{out}$ |
| *Twist-bending coupling* | $U_{tb} = \sum_{i=1}^{N-1} \frac{E_{tb_2}^i}{l_0^i}\left[m^i - m_0^i\right]\left[\kappa_2^i - \kappa_{2,0}^i\right]$ | 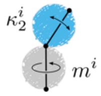 | $E_{tb_2}^i = E_{tb_2}^{in}$, if $i$ is odd<br>$E_{tb_2}^i = E_{tb_2}^{out}$, if $i$ is even | $E_{tb_2}^{in}$ $E_{tb_2}^{out}$ |
| *Lateral interactions* | $U_{lat} = \sum_{p,q}^{pairs} \frac{U_{lat}^0}{3}\left[1 - e^{-a_{lat}\left(r_{lat}^{pq} - r_{lat}^0\right)}\right]^2 + \frac{U_{LJ}^0}{3}\left[\frac{\sigma_{LJ}^0}{R_{lat}^{pq}}\right]^{12}$ | 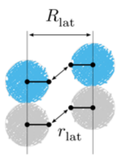 | | $U_{lat}^0$<br>$a_{lat}$<br>$r_{lat}^0$ |
| *Longitudinal interactions* | $U_{long} = \sum_{p,q}^{pairs} U_{long}^0\left[1 - e^{-a_{long}\left(r_{long}^{pq} - r_{long}^0\right)}\right]^2$ | 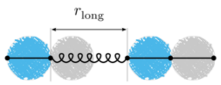 | | $U_{long}^0$<br>$a_{long}$<br>$r_{long}^0$ |

**Table S2.** Values of the model parameters used in this study. If longitudinal bonds are set to be breakable, the equilibrium edge lengths of α- and β-tubulin (top; marked with *) are replaced with the equilibrium bond distances of the Morse potentials (bottom; marked with *).

| Parameter | Units | Optimized or fixed from MD? | Value (GTP) | | Value (GDP) | |
|---|---|---|---|---|---|---|
| $E_s^{\alpha}$ | kJ/mol/nm | Optimized | 6061.6 | | 10492.3 | |
| $E_s^{\beta}$ | kJ/mol/nm | Optimized | 6868.7 | | 11397.3 | |
| $\lVert\mathbf{e}_0^{\alpha}\rVert$ | nm | Fixed | 3.79* | | 3.78* | |
| $\lVert\mathbf{e}_0^{\beta}\rVert$ | nm | Fixed | 4.17 | | 4.20 | |
| $E_t^{in}$ | kJ/mol/rad$^2$·nm | Optimized | 8036.8 | | 12931.5 | |
| $E_t^{out}$ | kJ/mol/rad$^2$·nm | Optimized | 7841.7 | | 12783.9 | |
| $m_0^{in}$ | rad | Fixed | -0.0147 | | -0.0103 | |
| $m_0^{out}$ | rad | Fixed | -0.0651 | | -0.1382 | |
| $E_{b_1}^{in}$ | kJ/mol·nm | Optimized | 4353.7 | | 3250.9 | |
| $E_{b_1}^{out}$ | kJ/mol·nm | Optimized | 6724.3 | | 13115.6 | |
| $E_{b_2}^{in}$ | kJ/mol·nm | Optimized | 16138.4 | | 18479.7 | |
| $E_{b_2}^{out}$ | kJ/mol·nm | Optimized | 16170.8 | | 17889.8 | |
| $\kappa_{1,0}^{in}$ | – | Fixed | -0.0889 | | -0.0451 | |
| $\kappa_{1,0}^{out}$ | – | Fixed | -0.3015 | | -0.2546 | |
| $\kappa_{2,0}^{in}$ | – | Fixed | -0.0040 | | -0.0122 | |
| $\kappa_{2,0}^{out}$ | – | Fixed | 0.0621 | | 0.0720 | |
| $E_{tb_2}^{in}$ | kJ/mol/rad·nm | Optimized | 8434.3 | | 9665.8 | |
| $E_{tb_2}^{out}$ | kJ/mol/rad·nm | Optimized | 8276.8 | | 9039.7 | |
| | | | *homo* | *seam* | *homo* | *seam* |
| $U_{lat}^0$ | kJ/mol | Optimized | 28.7 | 21.0 | 42.9 | 25.9 |
| $a_{lat}$ | nm$^{-1}$ | Optimized | 2.06 | 1.53 | 2.16 | 2.36 |
| $r_{lat}^0$ | nm | Fixed | 5.23 | 5.45 | 5.20 | 5.23 |
| $U_{long}^0$ | kJ/mol | Optimized | 58.3 | | 57.7 | |
| $a_{long}$ | nm$^{-1}$ | Optimized | 3.65 | | 4.81 | |
| $r_{long}^0$ | nm | Fixed | 4.42* | | 4.33* | |

**Movie S1 (separate file).** Movie showing the dynamic cluster formation and dissociation at a GTP-microtubule end. $L_{PF}$ = 6 dimers and $U_{lat}$ = 40 kJ/mol. The duration of the simulation is 200 μs sampled with a time step of 250 ns.

**Movie S2 (separate file).** Movie showing the dynamic cluster formation and dissociation at a GDP-microtubule end. $L_{PF}$ = 6 dimers and $U_{lat}$ = 40 kJ/mol. The duration of the simulation is 200 μs sampled with a time step of 250 ns.

### SI References


1. Igaev, M. & Grubmüller, H. Bending-torsional elasticity and energetics of the plus-end microtubule tip. *Proc. Natl. Acad. Sci. U. S. A.* **119**, e2115516119 (2022).
2. Jawed, M. K., Novelia, A. & O'Reilly, O. M. *A Primer on the Kinematics of Discrete Elastic Rods*. (Springer International Publishing, Cham, Switzerland, 2018).
